# Locally adapted oak populations along an elevation gradient display different molecular strategies to regulate bud phenology

**DOI:** 10.1101/2021.09.21.461210

**Authors:** Gregoire Le Provost, Céline Lalanne, Isabelle Lesur, Jean-Marc Louvet, Sylvain Delzon, Antoine Kremer, Karine Labadie, Jean-Marc Aury, Corinne Da Silva, Christophe Plomion

## Abstract

**Research conducted:** With the ongoing global warming, there are serious concerns about the persistence of locally adapted populations. Indeed, with the raising of temperature, the phenological cycle of tree species may be strongly affected since higher winter temperatures may have a negative impact on endodormancy release if chilling requirements are not fulfilled during winter and late frost in spring may expose trees if buds flush too early. Thus, Environmental gradients (showing continuous variations of environmental conditions) constitute a design of choice to analyze the effect of winter dormancy in locally adapted population.

**Methods:** In the present study, we used an elevation gradient in the Pyrenees to explore the gene expression network involved in dormancy regulation in natural populations of sessile oak locally adapted to temperature. Terminal buds were harvested during dormancy induction and release at different elevations. Then, gene expression was quantified using RNAseq and we used a likelihood ratio test to identify genes displaying significant dormancy, elevation or dormancy-by-elevation interaction effects.

**Key results:** Our results highlight molecular processes in locally adapted populations along this elevation cline, and made it possible to identify key dormancy-by-elevation responsive genes revealing that locally adapted populations have evolved distinct molecular strategies to adapt their bud phenology in response to environmental variation (i.e. temperature).

## Introduction

Despite extensive gene flow among populations (Kremer *et al*., 2012), forest trees demonstrate clear adaptation to local environmental conditions (Kawecki & Ebert, 2004; Savolainen *et al*., 2007; Aitken *et al*., 2008). There is indeed ample evidence of the efficiency of natural selection shown by the large body of literature on forest tree provenance tests (reviewed by (Alberto et al., 2013; Kremer et al., 2014). In almost any tree species for which provenance tests have been established, significant genetic variations between populations (i.e. differentiation) have been observed for fitness-related traits, among which bud phenology has taken a special place (White et al., 2007). Besides, these common garden experiments have clearly shown that phenotypic variation was structured along temperature and/or photoperiod gradients depending on the population of origin. The case of European white oaks, clearly illustrates the imprint of natural selection in shaping bud phenology variation and how forest tree populations became locally adapted in a short time span after the last glaciation (Ducousso *et al*., 1996; Vitasse *et al*., 2009b). Indeed, it has been shown that extant sessile and pedunculate oak populations stemming from the same or different source (refugial) of glacial origin, but growing today in different latitude exhibit strong phenotypic differentiation for the timing of leaf unfolding (Leroy *et al*., 2020) while these populations are not differentiated for neutral genetic markers (Zanetto & Kremer, 1995; Kremer *et al*., 2002) and the same holds true for populations distributed along elevation clines (Firmat *et al*., 2017; Leroy *et al*., 2020).

Environmental gradients (showing continuous variations of environmental conditions) produce spatially variable selective pressure on populations within the species’ range, therefore inducing genetic differentiation. The way in which genetic diversity is distributed across environmental gradients indicates how past selection has affected the genetic composition of populations and allows to identify parts of the genome that have undergone selection (Sork *et al*., 2016; Rellstab *et al*., 2016). In that regard, elevation gradients present several assets to discover genes that matter for adaptation: (i) phenotypic divergence between populations is generated by a predominant environmental driver -temperature - avoiding confounding effect with other potential drivers of adaptation, (ii) extensive gene flow between populations leaves the genome with narrow genomic windows where spatially varying selection is maintained and detectable, providing that genetic variation is studied throughout the genome, and (iii) populations have identical history avoiding confounding effect between demography and natural selection. Thus, elevation gradients have been extensively used to identify polymorphisms subjected to natural selection (Lobréaux & Miquel, 2020) including in forest trees (Leroy *et al*., 2020; Capblancq *et al*., 2020). As a complementary tool to the analysis of standing genetic variation for exploring genes of adaptive significance, gene expression data has also been a powerful approach to characterize functional genetic variation among populations (Mead *et al*., 2019). Comparison of expressional variations between locally adapted populations has been applied in many biological system and has allowed discovering genes of adaptive significance (reviewed by (Sork, 2018). More, combining allelic variation and gene expression data has also proved very powerful in identifying signature of adaptations in natural populations (Lasky *et al*., 2014; Cooper & Shaffer, 2021).

As stated previously, bud phenology constitutes a particularly poignant case study in the analysis of local adaptation of forest trees. In temperate region, the phenological cycle of the primary meristem is divided in two main phases: a growing period (i.e. from bud burst to bud set) when environmental conditions are favorable and a resting period also called dormancy, from bud set to bud burst when environmental conditions are unfavorable to stem elongation (Rohde & Bhalerao, 2007). Dormancy begins when growth cessation occurs in late summer as photoperiod and temperature decrease (Balandier *et al*., 1993); a phase called paradormancy or summer dormancy (Cline & Deppong, 1999). This first phase is followed by endodormancy. Endodormancy is initiated by lower temperature and shorter photoperiod in autumn (Heide, 2008). It is generally associated with a cold acclimation process (Tanino *et al*., 2010) and progressively induced until buds are unresponsive to growth in favorable conditions (Maurya & Bhalerao, 2017). Endodormancy is thus the deepest stage of dormancy and is released when chilling requirements are fulfilled (Luedeling *et al*., 2011), estimated around late January for oak populations in the Pyrenees (Dantec *et al*., 2014). In temperate regions, endodormancy generally starts during early autumn and reaches a peak in early winter according to the species considered and climatic conditions (Naor *et al*., 2003). Once chilling requirements are fulfilled, buds transition from endormancy to ecodormancy (Rohde & Bhalerao, 2007). During ecodormancy, bud development is triggered by forcing air temperatures in late winter and early spring. Overall, the timing of the phenological events observed in trees is mainly correlated with photoperiod (main factor triggering entry in endodormancy) and temperature (chilling temperature to break dormancy and warm temperature as the main factor triggering ecodormancy and bud swelling) (Menzel, 2002; Doi & Katano, 2008) and it is well known that predictive models of leaf unfolding which include both variables are currently the most accurate (Schaber & Badeck, 2003).

With the ongoing global warming (that is particularly pronounced in mountainous areas (Lenoir *et al*., 2008; Lenoir & Svenning, 2015) there are serious concerns about the persistence of locally adapted populations. Indeed, with the raising of temperature, the phenological cycle of tree species may be strongly affected since higher winter temperatures may have a negative impact on endodormancy release if chilling requirements are not fulfilled during winter (Dantec *et al*., 2014) and late frost in spring may expose trees if buds flush too early (Olsen *et al*., 2014). At the molecular level, many studies have been performed in trees to decipher the molecular mechanisms involved in dormancy induction and release. These studies revealed that dormancy regulation is mainly driven by molecular pathways related to phytohormones (Khalil-Ur-Rehman *et al*., 2017; Chao *et al*., 2017), carbohydrates (Min *et al*., 2017), temperature (Paul *et al*., 2014), photoperiod (Lesur *et al*., 2015a), oxygen species (Takemura *et al*., 2015), water (Lesur *et al*., 2015b) and cold acclimation.

In forest trees, most of the studies were conducted in poplars (Maurya & Bhalerao, 2017), oak (Ueno *et al*., 2013), spruce (Karlgren *et al*., 2013), chestnut (Santamaría *et al*., 2011) and beech (Lesur et al., 2015a). However, these studies were implemented using a restricted number of genotypes harvested only during endodormancy or ecodormancy and thus insufficient to provide a global view of internal and external drivers on dormancy induction and release. To our best knowledge, the most detailed study was carried out by Ruttink *et al*., (2007) in poplar. These authors identified key molecular pathways involved in bud formation and dormancy induction and release. They also identified a large set of genes commonly expressed during the growth-to-dormancy transitions in poplar apical buds, cambium, or Arabidopsis thaliana seeds, suggesting parallels in the underlying molecular mechanisms in various plant organs. Unfortunately, this study was conducted in controlled conditions for temperature and photoperiod. Moreover, none of the aforementioned studies have taken a population genetics perspective to account for natural variation in their ability to adapt their phenological cycle to varying external cues.

In the present study, we used an elevation gradient in the Pyrenees to explore the gene expression network involved in dormancy regulation in natural populations of sessile oak locally adapted to temperature. We addressed three main questions: Does gene expression in vegetative buds reflects divergent selection forces that have shaped population adaptation to elevation? Have locally adapted populations evolved different regulation strategies at endodormancy and ecodormancy stages? What gene functions have been targets of adaptation at the gene expression level? To this end, terminal buds were harvested during dormancy induction and release at different elevations. Since populations were continuously monitored for phenology over years we could account for their variability in bud burst date. Then, gene expression was quantified using RNAseq and we used a likelihood ratio test to identify genes displaying significant dormancy, elevation or dormancy-by-elevation interaction effects. Our results highlight molecular processes in locally adapted populations along this elevation cline, and made it possible to identify key dormancy-by-elevation responsive genes revealing that locally adapted populations have evolved distinct molecular strategies to adapt their bud phenology in response to environmental variation (i.e. temperature). Our findings provide a set of candidate genes at which allele frequency changes could be used in predictive models aiming at inferring the risk of mal-adaptation in assisted migration strategies in the frame of climate change (Rellstab *et al*., 2016).

## Material and methods

### Studied area and populations

We studied natural populations of sessile oaks (*Quercus petraea* (Matt.) Liebl.) located in two very close valleys (Ossau and Luz referred as O and L in the manuscript, respectively) of the northern slope of the Pyrenees Mountain, France (from 43°15′N, 00°44′W to 42°53′N, 00°06′E). In each valley, populations ranged from 100 to 1,600m a.s.l., representing a temperature gradient of 6.9°C between the lowest and highest elevated populations (Vitasse *et al*., 2009c). Functional traits related to phenology, morphology and physiology of these populations were extensively monitored *in situ* and *ex-situ* (i.e. in common gardens) during the past 15 years in order to quantify the respective contribution of phenotypic plasticity vs. genetic variation in shaping pattern of phenotypic variability (Alberto *et al*., 2011; Bresson *et al*., 2011; Dantec *et al*., 2014; Firmat *et al*., 2017). Using reciprocal transplant experiments, it was also shown that some of these populations are locally adapted (since some of these populations have the highest relative fitness at their home sites, and lower fitness in other sites), while other are not at their optimum (Vitasse *et al*., 2009a).

A general overview of the populations sampled in this study is provided in Supplementary Table 1 and Figure 1A.

**Figure 1:**
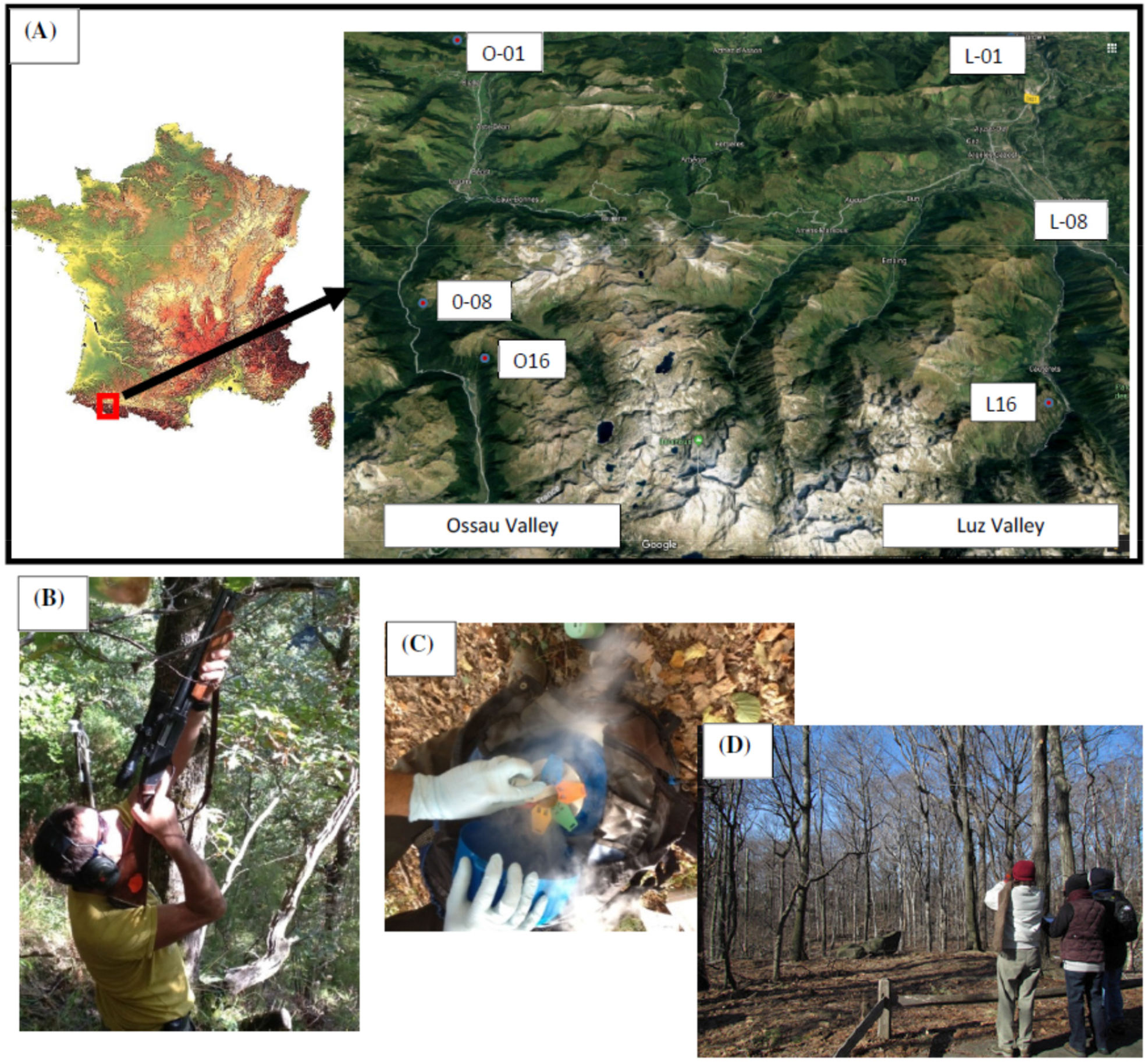
Sampling buds from sessil oak trees in the Pyrenees. A: Location of the three selected populations along two elevation clines (Ossau and Luz valleys). Population ID follows that of supplementary table 1. B: Shotgun to harvest terminal branches on three canopee, C: Bud storage in liquid nitrogen, D: Leaf unfolding observation with binoculars.

### Bud phenology monitoring and sampling

The timing of leaf unfolding was recorded for 8 years (between 2002 and 2014) in six populations on the same 20-34 adult trees per population, along the two elevation gradients. Leaf unfolding was monitored at 10-day intervals in each population, from March to June by the same two observers with binoculars (magnifying power 10, Figure 1E), about 15m away from the tree. At each visit, an assessment was made of the development state of each tree and leaf unfolding date after January 1^st^ (i.e. date of the year, DOY) was determined when leaves of 50% of its buds were fully unfolded following the method developed by (Vitasse *et al*., 2009b).

Based on their leaf unfolding timing, we selected three populations per valley representative of the intraspecific phenological variability of this elevation cline (Figure 2). These populations were located at 100, 800 and 1600m a.s.l. and exhibited substantial differences from one another in leaf unfolding timing (*ca* 50 days between the lowest and highest populations in elevation). Buds were sampled as follows: (i) a first campaign was performed in fall 2013 during the endodormancy period. Endodormancy is triggered by light and temperature but photoperiod plays a dominant role (Horvath *et al*., 2003) as demonstrated by short day experiments (Yang *et al*., 2021), thus trees were harvested during the same week. These samples corresponded to Endodormant buds, hereafter EndoD, (ii) a second campaign was performed in spring 2014 just before bud burst. As ecodormancy is mainly driven by forcing temperature (i.e. tree from high elevated populations flush later than those from lower elevation), we used the temporal series of leaf unfolding timing to determine the most relevant sampling dates at each elevation level. Thus, sampling was spanned over a period of 4 weeks from early March to early April to take into account this time-lapse in bud phenology (Supplementary Table 1). Samples corresponded to Ecodormancy, were referred hereafter as EcoD.

**Figure 2:**
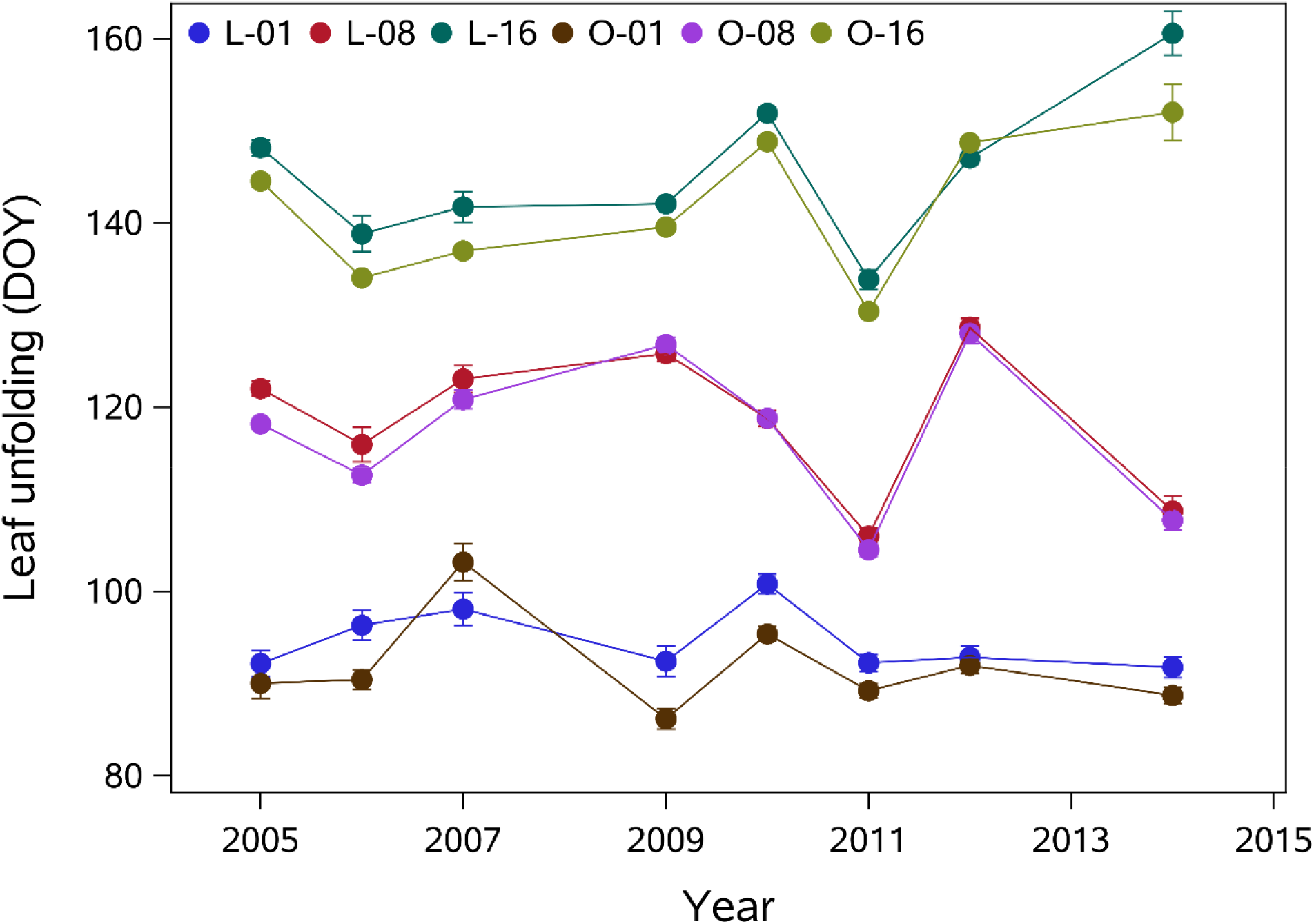
Mean date and standard error of leaf unfolding (y-axis, Date of the Year) for the three selected sessile oak populations in the Luz (L) and Ossau (O) valley at 100 (O-01, L-01), 800 (O-08, L-08) and 1600m (O-16, L-16) a.s.l.. Abbreviation of the populations follows that used in Figure 1.

For both developmental stages, buds were sampled on the same 20 trees in each population. We used a shotgun to sample terminal branches. Terminal buds were then harvested, immediately frozen in liquid nitrogen and stored at -80°C before subsequent analyses (Figure 1B and C).

### Hormones quantification

For hormone quantification, we used three pools of six individuals corresponding to three biological replicates for each population and dormancy stage (i.e. 36 samples in total). Buds were independently grinded in a fine power in liquid nitrogen before lyophilization, and then we mixed with an equimolar amount of the powder to constitute the pools. Phytohormones quantification were performed according to the procedure described by (Björklund *et al*., 2007)

### RNA extraction and sequencing

RNA extraction were independently performed for each individual and dormancy stage according to the procedure described by Le Provost *et al*., (2007). Residual genomic DNA was removed before purification using DNase RQ1 (Promega, Madisson, WI, USA) according to the manufacturer’s instruction. The quantity and the quality of each extract was then determined using an Agilent 2100 Bioanalyser (Agilent Technologies, Inc., Santa Clara, CA, USA). We generated 24 libraries corresponding to 2 dormancy phases x 3 populations x 2 Valleys x 2 biological replicates. For each biological replicate we pooled an equimolar amount of RNA extracted from 10 individual trees. Libraries were generated using the Illumina protocol (TruSeq Stranded mRNA Sample Prep Kit, Illumina, San Diego, CA, USA). Briefly, we selected mRNA form 2 µg of total RNA. mRNA were chemically fragmented and retro transcribed using random hexamer. After generating the second strand, the cDNA was 3’- adenylated, and we added the Illumina adapters. We amplified DNA fragments (with adapters) by PCR with Illumina adapter-specific primers. We quantified the libraries with a Qubit Fluorometer (Life Technologies, NY, USA). We estimated their size using the Agilent 2100 Bioanalyzer technology (Agilent). We then sequenced each library by 101 base-length read chemistry, in a paired-end flow cell, on an Illumina HiSeq2000 (Illumina).

### Trimming, mapping and identification of differentially expressed genes (DEG)

Cleaning and mapping were performed according to the procedure described in Le Provost *et al*., (2016). Briefly, we first removed low quality reads and kept sequences with a mean quality value higher than 20. Mapping of the reads on the 25,808 oak gene models published with the reference oak genome (Plomion *et al*., 2018) was performed with BWA (V.0.6.1) using default parameters. A minimum coverage of 100 reads was applied over the 24 analyzed samples resulting in 18K genes available for the analysis of gene expression. We first used the Median of Ratio Method (RMR) available in the DEseq2 method to normalize our data. This method is recommended for gene count comparisons between samples and for differential expression analysis (reviewed by (Anders & Huber, 2010).

Then, we detected differentially expressed genes (DEG) using the DESeq2 package with a P-value<0.01 after adjustment for multiple testing using a False Discovery Rate (FDR) of 5%. We also used a 2-fold change ratio to identify the most differentially expressed genes. We tested the effect of dormancy stage, elevation and their interaction effect using likelihood ratio tests. The dormancy and elevation effects were tested by comparing a statistical model without interaction (M1) with two reduced models for the dormancy (M2) and the elevation (M3) effects, respectively.

M1: Y_ijk_ = μ +D_i_ +E_j_ +ε_ijk_ where D_i_ is the dormancy stage (i= “Endodormancy” or “Ecodormancy”), and E_j_ the elevation effect (j= “low-100m”, “medium-800m” and “high-1,600m” elevation).

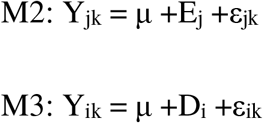

For the estimation of the interaction effect, we compared a complete model:

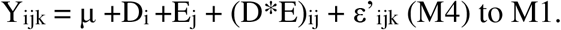

Three gene sets were thus generated (i) DEGs (geneset#1) corresponding to dormancy regulation (irrespective of elevation), (ii) DEGs (geneset#2) corresponding to elevation differences (irrespective of dormancy stage), and (ii) DEG (geneset#3) displaying a significant dormancy-by-elevation interaction. Annotation for each DEG was recovered from the pedunculate oak genome (Plomion *et al*., 2018). In the next sections, more emphasis will be placed on geneset#2 and #3, since they should include the key molecular players involved in local adaptation to temperature for the former, and reveal different strategies of locally adapted populations across bud phenological stages to cope with temperature variation for the latter.

DEGs from geneset#2 and #3 were analyzed using the EXPANDER software (Sharan *et al*., 2003) which cluster genes according to their expression profile using a Kmeans algorithm (Tavazoie *et al*., 1999). For both genesets, we set the number of clusters to 5 (k=5) to obtain the best homogeneity value within each cluster. Genes from each cluster were then used to perform independent gene set and subnetwork enrichment analysis (see next session).

### Gene set and subnetwork enrichment analysis

GO term enrichment analyses were first performed using the TopGO R package (Alexa, 2010). We used a corrected P-value lower than 0.05 to declare an ontology significantly enriched in our dataset.

Second, we performed subnetwork enrichment analysis using the Pathway Studio software (Pathway Studio®, Elsevier 2017) according to the procedure described in Le Provost *et al*., (2016).

### qPCR validation

For the elevation and the Dormancy-by-elevation effect, we selected six genes for qPCR validation. All the 12 genes analyzed with their associated effect are listed in Supplementary Table 2. qPCR quantification was performed with 1µg of total RNA according to the procedure described by Le Provost et al., (2012) on a Lightcycler® LC 384 (Roche Lifescience, Germany,

EU) with following standard parameters described in(Le Provost *et al*., 2016). Fluorescent data were normalized using two control genes (Qrob_P0530610 and Qrob_P0426000) and analyzed using STATQPCR (http://satqpcr.sophia.inra.fr/cgi/home.cgi). All the primer pairs used in this study were designed using the Primer3 software (Rozen & Skaletsky, 2000).

### Data accessibility

All the sequences generated for this publication have been deposited in the Europe Nucleotide Archive (https://www.ebi.ac.uk/ena) under accession <u>PRJEB17876</u>.

## Results

### Changes in phytohormone concentrations across elevations and dormancy stages

Endogenous concentration of three main phytohormones (indole-3-acetic acid (IAA), Abscicic acid (ABA) and Cytokinine (CTK)) was determined using three biological replicates for each dormancy stage, population and valley. First, we performed an ANOVA on the whole dataset to test if the two valleys could be considered as biological replicates. We used the following fixed-effects model: Yijk=µ+Di+Ej+Vk+ εi_ik_, where D_i_ is the dormancy stage (i= “Endodormancy” or “Ecodormancy”), and E_j_ the elevation (j= “low”, “medium” and “high” elevation) and V the valley effect (k= “Ossau” or “Luz”). Results are available in Supplementary Figure 1, panel A. No valley effect was significant. Thus, in a second analysis, we used the whole dataset, with the following fixed-effects model Yijk=µ+Di+Ej+Di*Ej+ ε_ij_, where D_i_ is the dormancy stage, and E_j_ the elevation effect to identify significant dormancy, elevation and dormancy-by-elevation interaction effects. Results are summarized in Supplementary Figure 1 Panel A.

For IAA and CTK a higher content was found in Ecodormancy whatever the elevation, while the opposite trend was found for ABA (Supplementary Figure1 Panel B). These results agree with those obtained by Yordanov et al., (2014) and Li et al., (2008) in ecodormant poplar and endodormant pear buds, respectively, where similar pattern of hormone dynamics along the winter period were observed.

A dormancy-by-elevation effect was identified for both IAA and ABA, respectively (Supplementary Figure1 Panel B). While IAA increases from low to high elevation in Endodormancy, it displays the reverse trend during ecodormancy. ABA increased from low to medium and high elevation in endodormancy but remained stable across the cline during ecodormancy.

### RNA-seq analysis and identification of DEGs

The main goal of our study was to identify genes involved in dormancy regulation (induction and release) and shape by natural selection, from the analysis of locally adapted populations to different elevations. To this end, we generated 24 cDNA libraries. A general overview of the libraries generated is shown in supplementary Table 3. Overall, more than 597 million reads were mapped onto the *Q. robur* reference genome. The mapping rates ranged from 66% to 82%.

We explore transcriptome-wide changes of our dataset with PCA. This exploratory analysis showed close clustering within biological replicates (including the “valley” effect), attesting the quality of the RNAseq data (Figure 3A). Besides, it showed that each dormancy stage clustered into two distinct groups (see the green and red dots separated along the first PCA axis in Figure 3).

**Figure 3:**
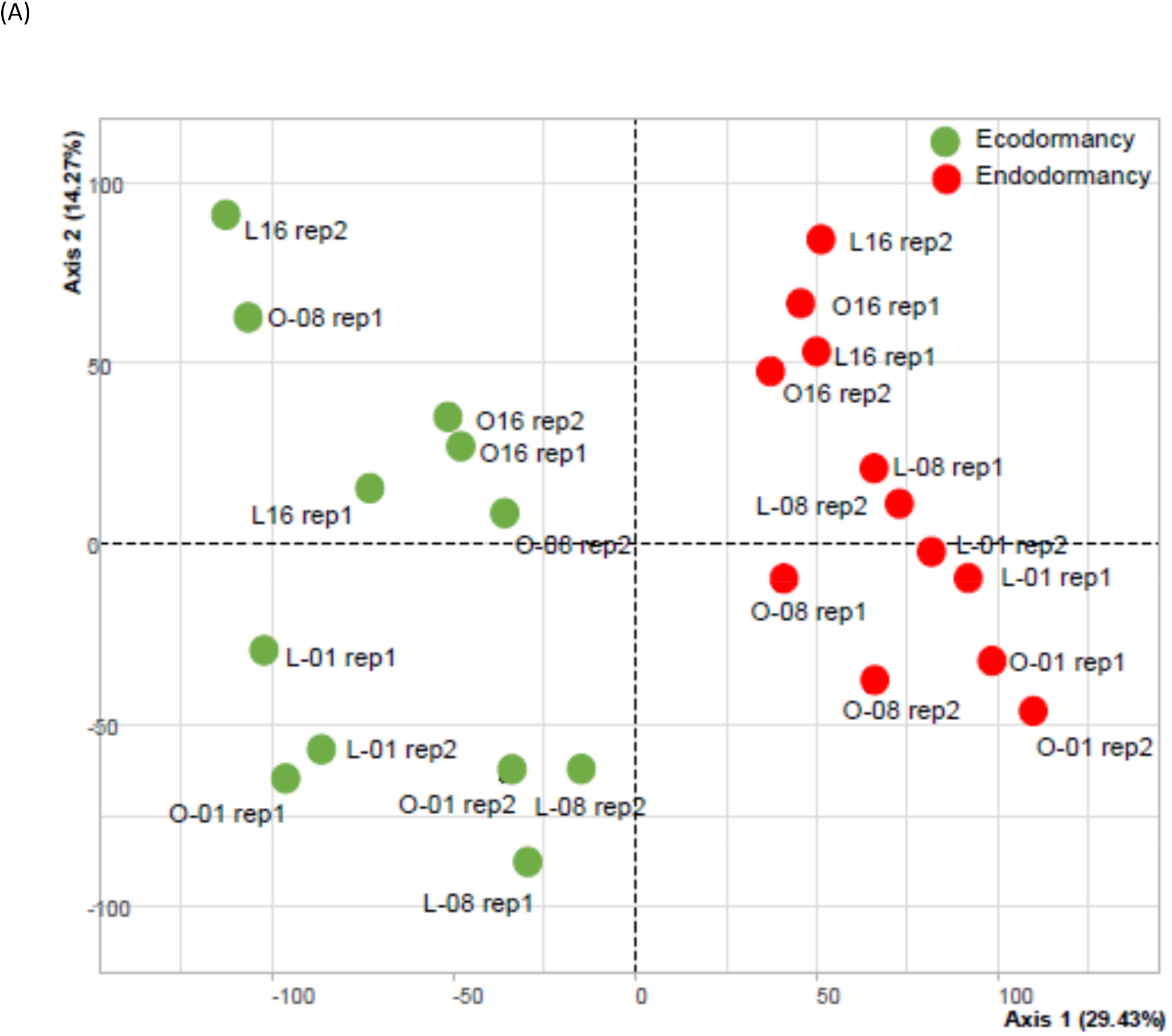
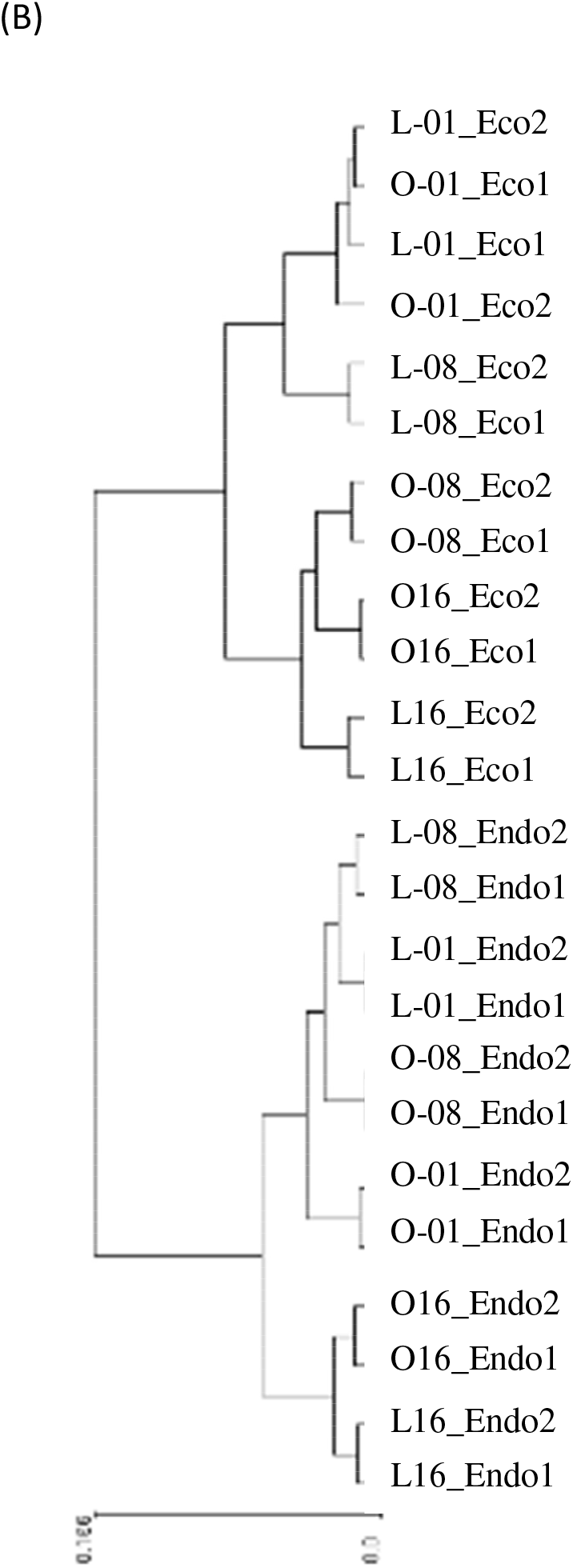
Evaluation of RNA-seq data quality using PCA(A) and clustering analysis (B) (using the “complete” method of the expender software) among the replicates of the different dormancy stages and elevations. For PCA red and green dots represent endo- and eco-dormancy samples, respectively. The sample names follow that of Supplementary Table 1.

The adjusted data were also evaluated with a clustering approach using the complete method available in the Expender software. This analysis confirmed the high level of reproducibility of the RNAseq data and revealed for each treatment analyzed in this study that the biological replicates and dormancy stages were grouped in a same cluster (Figure 3B). Both the PCA (along the second PCA axis) and clustering also revealed the presence of interaction between dormancy and elevation. A total of 17,676 genes remains following quality based filtering.

A preliminary analysis was performed to test that the two valleys could be considered as two biological replicates for the RNAseq data. For each dormancy stage we compared gene expression level between pairs of populations samples at the same elevation (i.e. O-01 vs. L-01, O-08 vs. L-08, O-16 vs. L-16) resulting in six different gene sets (2 dormancy stages * 3 elevations). Results are summarized in Supplementary Table 4. Whatever the dormancy stage considered, we identified very few genes differentially regulated between the two valleys and reaching a maximum of 1.2% of the 17,676 genes between O-08 vs. L-08 at endodormancy. These results confirmed that the two valleys could be considered as biological replicates. Increasing the number of biological replicates (from 2 to 4) contributed to reduce the false-positive rate and thus to distinguish noise to a true biological signal in our dataset.

We then used the likelihood ratio tests to identify genes displaying significant dormancy, elevation and interaction effects. Genes exhibiting differential expression by at least two-fold with the corrected P-value (<0.01), and adjusting with false discovery rate (q-value) < 0.05 were deemed significantly differentially expressed. Results of the differential expression analysis are shown in Figure 4, panel A. The overlap between the three effects is shown in Figure 4, panel B. It should be noticed that a very low number of genes (25) identified in this study displayed the three effects simultaneously. Overall 2,084, 1,089 and 635 genes displayed a significant Dormancy, Elevation or Dormancy-by-elevation effect, respectively. 397 genes were regulated both by dormancy and elevation and we identified 203 genes displaying significant dormancy and dormancy-by elevation interaction effects (Figure 4 Panel B and C). Finally, 80 genes were significant both for elevation and dormancy-by-elevation interaction effects.

**Figure 4:**
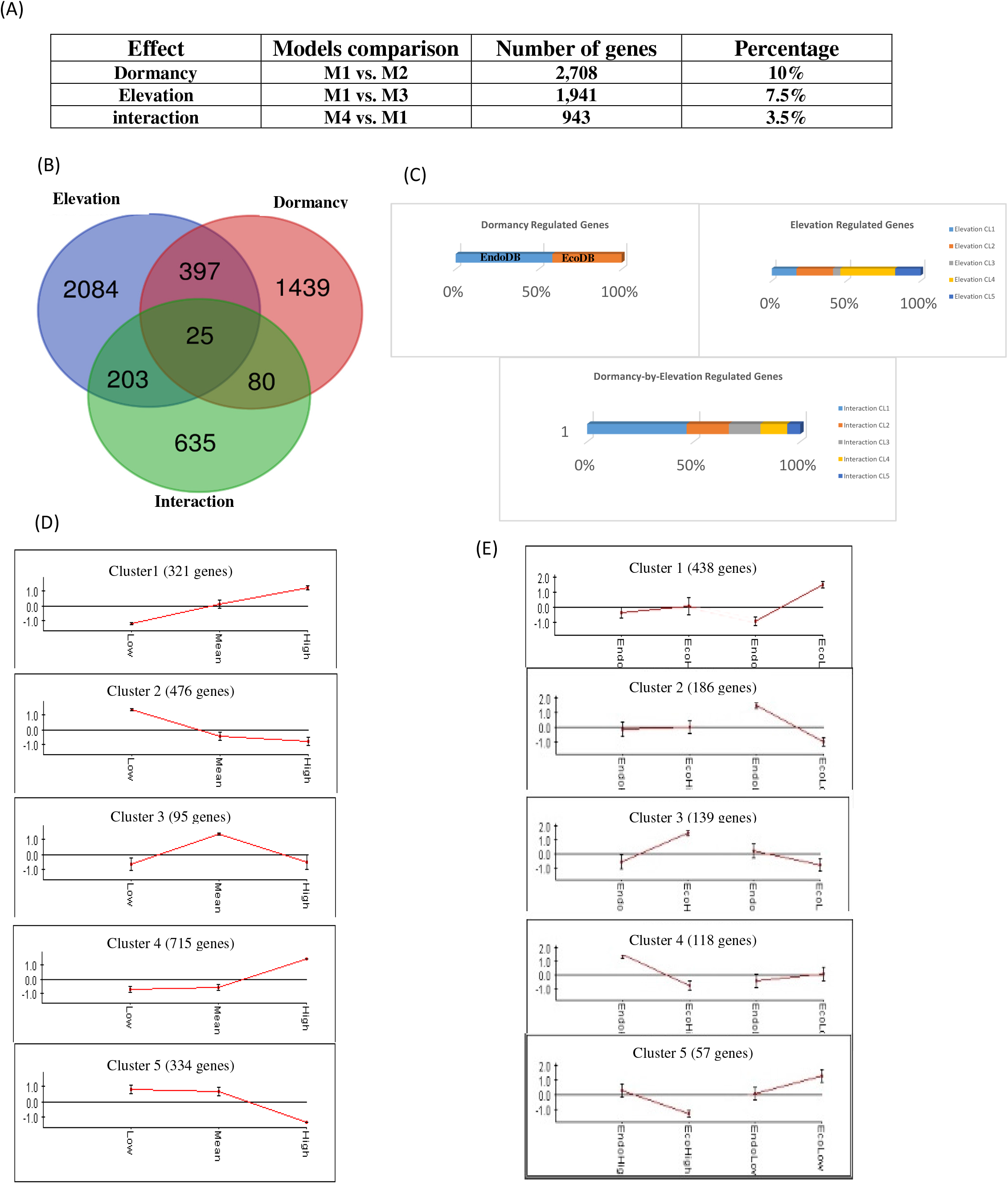
Illustration of the main results obtained from the DEGs analysis. Panel (A): Summary of the differential expression analysis. Results are shown for an adjusted P-value<0.01 and a fold-change ratio>2. The percentage indicated in the last column is given considering the 25,808 gene models found in the pedunculate oak genome. Panel (B): Venn-diagram showing the number of genes displaying main (Dormancy, Elevation) and Interaction effects or a combination of them. Panel (C): Bar Chart illustrating the number of genes identified per effect and their associated clusters. Panel (D) Cluster identified for the elevation responsive genes and Panel (E) clusters identified for the dormancy by elevation responsive genes.

### Genes regulated during dormancy induction and release (Geneset#1)

By comparing model M1 and M2, using the threshold presented above, we identified 2,709 genes differentially regulated across these two dormancy stages (Figure 4, panel A). For this first gene set, we mostly identified quantitative variations (*i*.*e*. all the genes displaying a differential expression were covered by reads originated both from Endodormancy and Ecodormancy libraries). We identified a single gene (Qrob_P0090160.2, encoding a riboflavin synthase) covered by reads from the Ecodormancy samples only. Detailed results are available in Supplementary File 1. Gene set enrichment analysis was performed using the TopGo Package for this first gene set. Regarding the Molecular function ontology, the most enriched GO terms were related to “catalytic activity”, “heme binding”, “oxygen oxidoreductase activity”, “tetrapyrrole binding” and “oxidoreductase activity” for endodormancy responsive-genes, and “structural constituent of ribosome”, “structural molecule activity”, “rRNA binding”, “urea transmembrane transporter activity” and “copper ion binding” for ecodormancy responsive-genes. Regarding the Molecular Function ontology, we identified terms related to “catalytic activity”, “heme binding”, “hydroquinone:oxygen oxidoreductase activity”, “tetrapyrrole binding” and “oxidoreductase activity” for the former stage, while terms related to “structural constituent of ribosome”, “structural molecule activity”, “rRNA binding”, “urea transmembrane transporter activity” and “copper ion binding” were identified for the latter. We performed also subnetwork enrichment analysis independently for genes regulated during endodormancy and ecodormancy. While we identified biological processes related to cold tolerance, response to dehydration, freezing tolerance and drought tolerance in Endormant buds, very different biological processes related to meristem identity, flower development and floral organ identity were identified in Ecodormant buds. These results agree to that observed by Ueno *et al*., 2013 during dormancy induction and release in a previous study in oaks, thus validating our experimental design and statistical model in finding dormancy regulating genes and opening encouraging perspectives to discover elevation and interaction related genes (see next sections).

### Elevation regulated genes (Geneset#2)

By comparing models M1 and M3, we identified 1,941 genes regulated by elevation (Figure 4, panel A and Supplementary File 2)). This gene set may encompass mostly genes involved in adaptation to temperature. We cluster these genes according to their expression profile. We set the number of clusters to 5 to obtain an overall homogeneity close to 0.98. Cluster#1 included 321 genes gradually up-regulated by elevation; conversely cluster#2 contained 476 genes showing the reverse trend, while cluster#3 comprised 95 genes upregulated at the intermediate elevation level. Cluster#4 and #5 were strongly differentiated showing contrasted gene expression patterns. Cluster#4 contained 715 genes over-expressed at the highest elevations, while cluster#5 encompassed 334 genes down-regulated at highest elevation (Figure 4, panel D and supplementary File 2).

Gene set enrichment analysis was then performed for each cluster (Supplementary File 2). Regarding the Molecular Function ontology, we identified for: (i) cluster #1 enriched terms related to “quercetin 7-O-glucosyltransferase activity”, “UDP-glucosyltransferase activity”, “quercetin 3-O-glucosyltransferase activity”, “raffinose alpha-galactosidase activity” and “alpha-galactosidase activity”, (ii) cluster #2 enriched terms related to “alcohol dehydrogenase (NAD) activity”, “benzoic acid glucosyltransferase activity”, “polysaccharide binding”, “pattern binding” and “quercetin 7-O-glucosyltransferase activity, (iii) cluster #3, enriched terms related to “adenyl ribonucleotide binding”, “adenyl nucleotide binding”, “purine ribonucleotide binding”, “purine nucleotide binding” and “ribonucleotide binding”, (iv) cluster#4 enriched terms related to “transcription factor activity”, “nucleic acid binding”,” calcium binding”, “protein serine/threonine phosphatase activity” and “phosphatase activity” “, and (v) for cluster#5, no statistical enrichment was identified.

Subnetwork enrichment analysis was performed for each of the five clusters (Supplementary File 2). We identified hubs related to leaf size, root growth and pollen development for genes belonging to cluster #1, associated with senescence, cell death and cell division for cluster #2, root development, root growth and flower development for cluster#3, internode patterning, fatty acid omega oxidation and pedicel development for cluster #4, heat tolerance for cluster #5.

### Genes displaying a significant dormancy-by-elevation interaction effect

Using the likelihood ratio test comparing model M4 and M1, we identified 943 genes displaying a significant dormancy-by-elevation interaction effect (Figure 4, panel A and Supplementary File 3). Interestingly, most of these genes displayed only this effect (635 out of 943 genes, Figure 4). Only 25 genes displayed the three main effects simultaneously. The remaining genes (*i*.*e*. 283 genes) were significant for two effects: 203 genes with dormancy and 80 with elevation. In order to cluster genes according to their expression profile, the Kmeans was also applied to this gene set. A stable homogeneity value of approximately 0.91 was obtained for a value set to five allowing us to produce five different clusters (Figure 4, panel E and Supplementary File 3). Cluster #3 (139 genes) and cluster #2 (186 genes) displayed a similar expression profile characterized by a singular pattern where genes are upregulated during ecodormancy at high elevation and during endodormancy at low elevation. Cluster #1 (438 genes), cluster #5 (57 genes) and cluster #4 (118 genes) encompassed genes upregulated in ecodormancy at low elevation, while their expression level was either stable or down-regulated in the same developmental stage at high elevation.

Regarding the gene set enrichment analysis, a single GO term related to “regulation of biological process” was enriched in cluster #3, while GO terms related to “homeostatic process”, “regulation of shoot system development” and “signal transduction” were identified in cluster #2. Similarly, a single GO term related to “developmental process involved in reproduction” was identified in cluster #5, while terms related to “cell cycle”, “negative regulation of biological process” and “regulation of cellular process” and terms related to “cellular lipid metabolic process”, “lipid biosynthetic process” and “lipid metabolic process” were enriched in cluster #1 and cluster #4, respectively.

Subnetwork enrichment analysis is also shown for each cluster in supplementary File 3. For cluster #2 and cluster #3, characterized by genes upregulated during ecodormancy for high elevation populations, we identified significant biological processes related to stress response in these two clusters, namely: genes related to “salinity”, “plant defense” and “plant development” in cluster #2, and genes involved in “salinity response”, “detoxification” and “cell elongation” in cluster #3. For cluster #1, #4 and #5, characterized by genes upregulated in ecodormant buds for low elevation populations, we mostly identified biological processes related to meristem functioning. Important hubs related to “plant growth”, “mRNA splicing” and “transcription activation” were identified in cluster #1. Regarding the two other clusters (#4 and #5) we identified hubs related to “plant growth”, “flower development” and “seed germination” for the former, and “cell elongation”, “developmental process” and “flower development” for the latter.

### qPCR validation

A total of 12 genes were selected to validate their expression profile by real time quantitative PCR. Among these genes, two displayed both multi-banding pattern on agarose gel analysis or failed PCR amplification and were removed from the analysis. For the other 10 genes, PCR efficiency ranged from 95 to 110 %, which is in accordance with Taq polymerase activity (Supplementary Table 2). The expression patterns of the tested genes revealed by qPCR were similar to those observed in RNAseq-data (supplementary Figure 2) validating our finding using RNA-seq data.

## Discussion

The main goal of this study was to provide new insights on key molecular pathways involved in dormancy regulation from the analysis of gene expression in populations locally adapted along an elevation cline. To reach this objective, we considered the two main dormancy phases endodormancy *vs*. ecodormancy and performed differential gene expression analysis from terminal vegetative buds using an RNAseq approach. We identified genes displaying significant expression variations associated with dormancy, elevation and dormancy-by-elevation interaction. While variation of dormancy related traits have been rather well investigated using latitudinal cline, where both photoperiod and temperature vary and influence local adaptation (Ingvarsson *et al*., 2006; Hall *et al*., 2007), elevation clines located on a restricted area present strong assets to disentangle these two environmental factors and offer a unique opportunity to analyze the effect of temperature independently from the photoperiod (Vitasse *et al*., 2009b). Moreover, to our best knowledge elevation clines have never been used to analyze the molecular mechanisms involved in bud dormancy regulation in a forest tree species. It is also well known that an increasing of temperature in the coming years will have a strong effect on the phenological cycle of perennials species (Tanino *et al*., 2010; Olsen *et al*., 2014). Therefore, we will emphasized our discussion on the genes regulated by elevation (i.e. temperature-responsive genes) and on those displaying a significant dormancy-by-elevation interaction effect, because these two sets of genes may encompass genes that matter for the adaptation of forest trees.

### (i) Developmental molecular plasticity underlying the shift between endo and ecodormancy

We quickly summarized here the main finding of our study that align on what is known from the literature on bud dormancy regulation.

Three phytohormones were first quantified (IAA, ABA and cytokinines) to phenotypically characterize the samples. As found in fruit trees (Li *et al*., (2018)) ABA content was found to be higher in endodormant buds. Several authors also reported that ABA is involved in the photoperiodic control of growth and potentially in growth cessation induced by short day (Maurya & Bhalerao, 2017). Conversely, the concentration of IAA and cytokinines were higher in ecodormant buds. IAA concentration is known to increase during cambium activity reactivation (Li *et al*., 2013) and cytokinines are activated during budburst (Dierck *et al*., 2016). Altogether, these results suggest that the higher contents of these two phytohormons in ecodormancy is correlated to the restart of the mitotic activity in the buds of the sampled trees. At the transcriptional level, we identified two biological networks corresponding to gene set#1 related to either endodormancy or ecodormancy (Supplementary File 1). As expected, important hubs related to cold tolerance, freezing tolerance, drought tolerance and defense response were identified in the former, while biological hubs related to meristem functioning (meristem identity, flower development, floral organ identity….) were enriched in the latter. Similar results were previously found in oak (Ueno *et al*., (2013)) and beech (Lesur *et al*., (2015a)) but the breath of our dataset extended our finding to much more genes as well as low expressed transcripts (transcription factors) compared to these studies. In endodormancy, the upregulation of molecular mechanisms involved in dehydration and cold tolerance is consistent with the fact that cold tolerant dehydrated tissues are less prone to ice formation and can therefore better cope with low temperature observed in winter (Horvath, 2010). In ecodormancy, central hubs related to cell activity and meristem functioning were identified reflecting the restart of the cellular machinery which is essential to bud burst when environmental conditions become favorable (Zhong *et al*., 2013; Shim *et al*., 2014).

### (ii) Environmental molecular plasticity to temperature

In mainland France, whatever modeled climatic scenario is, the mean temperature might increase from 0.6°C (North of France) to 1.3°C (South of France) by 2050 and an even greater temperature contrast is expected in the mountainous areas (Allamano *et al*., 2009). This increase in temperature will strongly influence the distribution of forest ecosystems (Tylianakis *et al*., 2018). Thus the elevation cline used in this study with a temperature lapse rate of 6.9°C constitute a unique opportunity to analyze the role of gene expression in local adaptation to temperature.

#### (a) Populations from low and medium elevations

Two clusters mainly encompass genes regulated at low and/or medium elevation (Cluster #2 and #5, respectively, Supplementary File 2, Figure 4, panel D). Population from these two clusters are characterized by an extended growing season associated to an earliest budburst date (Figure 2). This typical phenology may be explained by the main molecular mechanisms identified in our subnetwork enrichment analysis.

First, we identified an important number of genes in Cluster #2 related to the inhibition of cell death suggesting an extended cell life duration allowing the tree to extend its growing period. Indeed, half of the genes related to the cell death biological process are involved in cell death inhibition (CAS1, BON3, GLIP1, TOPII, T3G21.18 and F5O11.34). For example, we identified a CAS1 gene encoding a clycloartenol synthase 1 (4.2 fold-change ratio -FC-between low and high elevation populations). Kim *et al*., (2010) using downregulated Arabidopsis mutants reported that transgenic lines were characterized by a cell death phenotype suggesting a key role of the CAS1 gene in the inhibition of cell death. I the same way, BON3 (3.8 FC), a gene similar to a copine-like protein, has been reported to be involved in the inhibition of cell death (Kim *et al*., (2017)). A TOPII gene (2.24 FC) similar to a Topoisomerase protein was found to be downregulated in rice, in relation to a delayed cell death (Yang *et al*., 2012b). We also identified 15 genes belonging to the senescence biological process, including five genes (MYB62 (2.45 FC), AGL15 (4.35 FC), LOX2 (2.15 FC), ABI5 (2.81 FC) and SAG13 (2.23 FC)) involved in the inhibition of the senescence suggesting a later leaf fall in the population from lower elevation. As an example, the MYB62 gene which encodes a R2R3 MYB transcription factor is known to be involved in apical dominance, delayed flowering time and late senescence in sunflower (Moschen *et al*., 2014). Besides, the AGL15 gene encoding an Agamaous like protein has been shown to have a clear role in a delayed senescence in *Arabidopsis thaliana* (Fang & Fernandez, 2002).

Finally, 9 and 23 genes were found to be involved in the cell division and root growth, respectively, suggesting a higher meristematic activity in population from lower elevation allowing trees to flush earlier and to extend their growing period. Among those, ERF3 is similar to an Ethylene responsive factor which stimulates cell division in the root tip in rice (Zhao *et al*., 2015). Additionally, MYB12 is known to promote cell division and elongation in Arabidopsis (Tan *et al*., 2019).

A single biological process related to “Heat tolerance” was enriched in cluster #5 (Supplementary File 2). In this hub, all the genes are involved in thermotolerance acquisition. In higher plant, this mechanism allows plants to cope with heat stress and to maintain their productivity under unfavorable environmental conditions (reviewed by Burke, 2001). We hypothesized that a better thermotolerance acquisition in population either from lower elevation is an efficient strategy to cope with stressfull conditions that are more frequently encountered in lower altitude populations that display an extended growing period. Among these, two genes retained our attention: AGO1 (FC=2.26) and TOR (FC=2.55). AGO1 encodes an RNA Slice protein known to be essential for the maintenance of the acquired thermotolerance (Stief *et al*., 2014). Besides, the downregulation of TOR (similar to a phosphatidylinositol 3-kinase family protein TOR, AT1G50030) in the shoot apical meristem of *Arabidopsis thaliana* causes weaker thermotolerance (Sharma *et al*., (2019)).

#### (b) Population from high elevation

Two other clusters (#1 and #4) comprised genes upregulated at higher elevation (Supplementary File 2, Figure 4, panel D). Oak populations from high elevation flush latter and their vegetative phase, much shorter, occurs when environmental conditions are favorable (Vitasse *et al*., 2009b). Moreover, their leaf size is smaller compared with those of low elevation population (Bresson *et al*., 2011). The molecular mechanisms we identified in the subnetwork enrichment analysis may explain these observations. Indeed, biological processes related to “leaf size”, “pollen development”, “root growth”, “internode patterning” and “pedicel development” were found in cluster#1 and #4. In the cluster #1, ten genes are involved in the leaf size biological process. Among those, three genes (POL (FC=2.47 between high and Low elevation population), T1M15.220 (FC=2.21) and GAI FC=2.63) are implicated in the reduction of leaf size while the other are essential for a normal leaf development. POL (encoding a Protein phosphatase 2C) is known to be involved in the reduction of leaf size in Arabidopsis (DeYoung *et al*., 2006). T1M15.220 (similar to a SAUR protein) is also known to be associated with a smaller leave size in plant (reviewed by Ren & Gray, 2015). Finally, overexpression GAI (GRAS family transcription factor) leads to a smaller leaf size in petunia (Gargul *et al*., 2017).

In cluster#1, we also found 16 and 7 genes related to root growth and meristem size, respectively. Most of the genes identified in this cluster are essential for meristem functioning which illustrates once again the ability of the high elevation populations to produce leaves over a short growing period. As an example, the BIG gene (Auxin transporter protein, FC=2.6) is known to be essential for cell proliferation in Arabidopsis roots and shoot meristems in response to sucrose and glucose (Ayala-Rodríguez *et al*., 2017). Also, RPK2 (similar to a receptor-like kinase RPK2, FC=2.36) is known to be involved in the regulation of the meristem size by controlling cell proliferation (Racolta *et al*., 2018).

Eleven genes belonging to the “pollen development” biological process were also identified suggesting that the cellular machinery must be activated in the population of high elevation to flush and flower rapidly when environmental conditions become favorable. Among them we identified three GSL genes (GSL1 (FC=2.32), GSL8 (FC=2.49) and GSL10 (FC=2.62)) encoding callose synthase. In *Arabidopsis thaliana*, callose synthase has been reported to play a key role during pollen development (Zhu *et al*., 2018). We also identified one RGL genes (RGL2 (FC=2.77) and RGL1 (FC=3.47) encoding DELLA protein. In Arabidopsis, Fleet & Sun, (2005) reported that DELLA gene repressed petal and pollen development as well as anther elongation suggesting a possible role of these genes in the delayed flowering time observed in this population.

Finally, only one biological process related to “internode patterning” containing three genes (KNAT1 (FC=2.63), STM (FC=4) and BLH8 (FC=2.43)) was identified in cluster#4 (Supplementary File 2). Among these genes, two (STM and BLH8) control meristem formation and/or maintenance, organ morphogenesis, organ position, and several aspects of the reproductive phase in Arabidopsis thaliana, suggesting a key role of these genes in meristem functioning (Hamant & Pautot, 2010).

In conclusion, our results highlight that main molecular mechanisms associated with genes differentially expressed at low and high elevations may explain the typical phenology and adaptive features to temperature of these locally adapted populations. Indeed, at low elevation, populations display a higher basal expression of gene involved in the inhibition of cell death and senescence, thermotolerance acquisition and in the maintenance of cell division activity allowing the trees to flush earlier when environmental conditions are favorable (Figure 5). On the other hand, populations from high elevation are characterized by overexpression of genes involved in leaf size reduction, meristem functioning and a delayed flowering time due to the shorter growing season observed in these trees (Figure 5).

**Figure 5:**
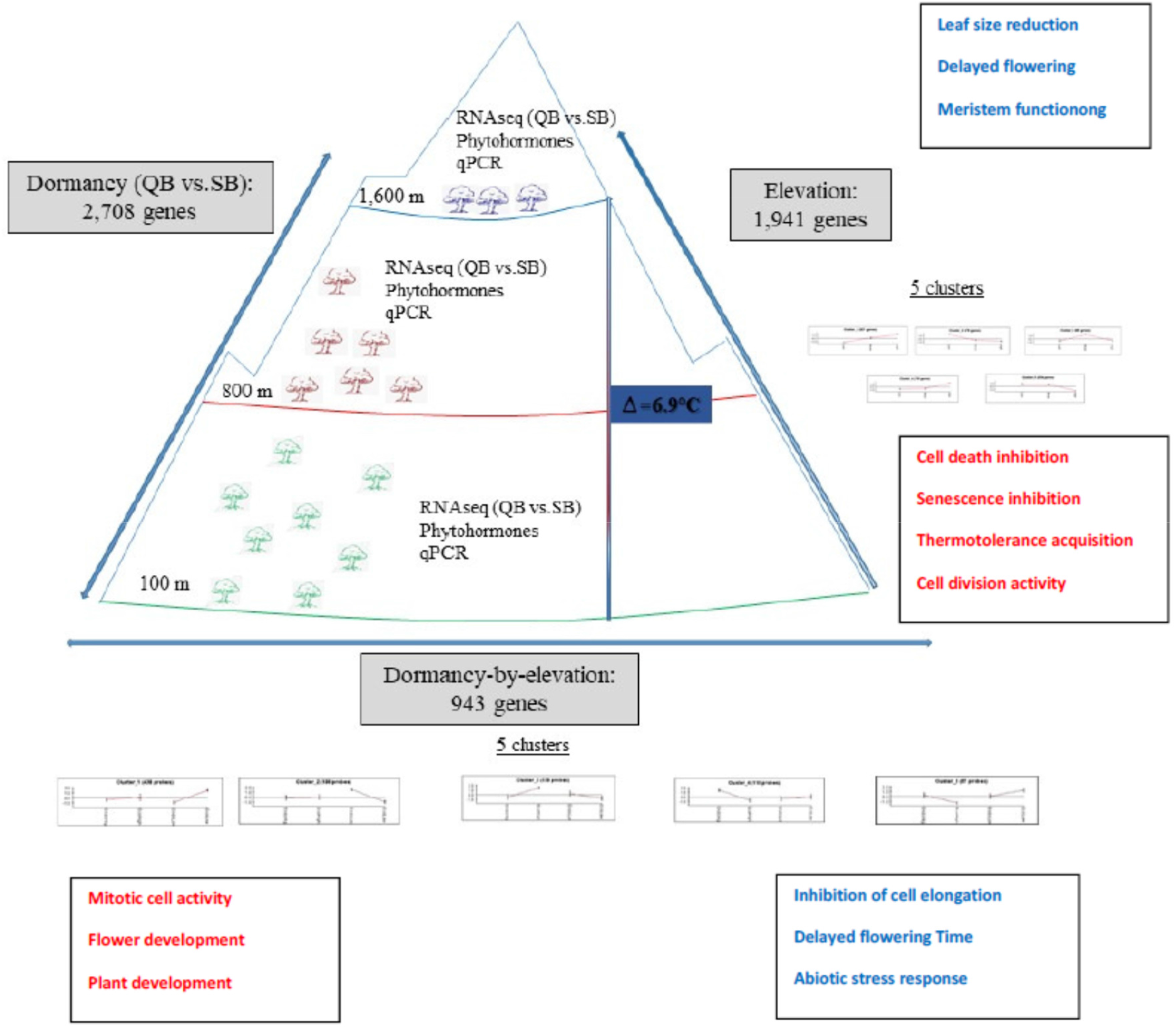
Overview of the results obtained in this study. Main molecular functions associated to elevation and dormancy-by-elevation interaction effect are indicated in color. Red color was used for population from low elevation, Blue color for population from high elevation.

### (iii) dormancy-by-elevation responsive genes reveal distinct molecular strategies of locally adapted populations to cope with temperature variation

The five clusters identified using the Kmeans approach were merged in two mains groups. A group of genes upregulated in ecodormant buds at high elevation (clusters #2 and #3, Figure 4 panel E) and a group of genes upregulated in ecodormant buds at low elevation (clusters #1, #4 and #5).

Gene pool from high elevation correspond to late flushing populations (Vitasse *et al*., 2009a). They have to cope with severe environmental conditions. A 6.9°C temperature gradient has been reported along the studied elevation cline. Once chilling recruitment is fullfilled, buds in the high elevation populations enter the ecodormancy phase. But they still have to cope with late damage frost currently observed at this elevation during early spring. The main molecular mechanisms identified in clusters #2 and #3 (*i*.*e*. flowering time, plant development, salinity response and detoxification (*i*.*e* cold tolerance)) may explain the typical bud phenology observed in these high elevation populations.

On the other hand, populations at low elevation comprised early flushing trees characterized by a longer growing period (Vitasse *et al*., 2009a). In general, at ecodormancy, populations from low elevation do not have to cope with late damage frost. We thus assume that the main biological processes found in clusters #1, #4 and #5 (cell activity, meristem functioning and flower development) prepare the buds to flush as soon as the heat requirement is fulfilled, thus explaining the longer growing period observed in these populations.

Given this, the analysis of genes showing an elevation x dormancy phase interaction effect provide clues about the following complementary questions: how populations at high elevation cope with late frost damage at ecodormancy? And which molecular mechanisms may explain the early flushing date observed in populations at low elevation.

#### (a) how do population from high elevation cope with adverse environmental conditions in ecodormancy to avoid late frost damage (cluster#2 and #3)

The network enrichment analysis performed for clusters #2 and #3 (Figure 4, panel E) identified several genes involved in the inhibition of cell elongation and/or in a delayed flowering time but also in the response to abiotic stresses. Regarding the flowering time biological process, we identified the ABF4 (bzip transcription factor) known to be involved in the inhibition of flowering time in Arabidopsis (Wang *et al*., 2013a). The WRKY71 transcription factor, also found in cluster#2, affects directly the flowering time of plants by regulating the Constance and Flowering locus genes (Jue et al., (2018)). The CRY1 gene (Cryptochrome 1 Apoprotein) is involved in the regulation of photoperiodic flowering and in the inhibition of cell elongation (He *et al*., 2015). MFB16.6 (encoding SPL13 protein) was also found to be upregulated during ecodormancy at high elevation. In Medicago, Gao *et al*., (2018) reported a potential role of this gene in a delayed flowering time. We also identified many genes involved in the regulation of cell elongation and generally in its inhibition suggesting a potential role of these genes in slowing meristem development. Among those, SWN and At3g51890 encode a swinger and the CLC3 protein, respectively. Footitt *et al*., (2015) reported a maximum expression of the SWN gene coincident with the dormancy peak in Arabidopsis suggesting a key role of this gene in dormancy release. Wang *et al*., (2013b) identified the CLC3 gene as a key molecular player involved both in basipetal transport and sensitivity and distribution of auxin. However, as CLC3 is mainly involved in auxin transport, it is difficult to relate its expression level to auxin content in our biological samples.

Moreover, we identified in cluster#3 genes potentially involved in the acceleration of cell elongation or in the activation of flowering that may explain the need to grow and achieve reproduction in a shorter period of time in these populations. BZR1 (encoding a brassinosteroid signaling positive regulator) is known to induce cell elongation at elevated temperature under the combined effect of auxin and brassinosteroid (Bouré *et al*., 2019). ROT3 encoding a cytochrome P-450 gene known to be involved in the leaf expansion rate in response to brassinosteroid (Bancoş *et al*., 2002). We identified also a CIB1 gene similar similar to a Cryptochrome-Interacting-Basic-Helix-Loop-Helix protein. Liu *et al*., (2018) showed that cryptochrome are blue light receptors that mediate light responses in plants and animals and activate flowering once environmental conditions are favorable (temperature and photoperiod).

Finally, we identified several genes related to “salinity response”, “plant defense” and ‘detoxication”. This is not surprising since this should allow the bud to stay dehydrated during ecodormancy to better cope with the low temperature frequently observed during early spring in populations from high elevation. In plant, several authors reported also that detoxication processes are essential to acquire cold tolerance (Koehler *et al*., 2012; Yang *et al*., 2012a). As example, we identified an APX3 encoding an ascorbate peroxidase involved in ROS detoxication. Wang & Li, (2006) reported that APX gene are overexpressed during cold acclimation. We also found an OSSA1 gene similar to a cytosolic O-acetylserine(thiol)lyase known to be involved in ROS detoxication in plant (Shirzadian-Khorramabad *et al*., 2010). Two GSTU genes (GSTU17 and GSTU9) similar to glutathione transferase protein were also identified. Fujino & Matsuda, (2009) showed that GSUT gene contribute to cold tolerance in rice. Regarding the PR4 gene which encodes a pathogenesis related protein, Cabello *et al*., (2012) reported that its induction is responsible for cold tolerance in Arabidopsis.

#### (b) Which molecular mechanisms are related to the early flushing date observed in population from low elevation

Our subnetwork analysis performed on clusters #1, #4 and #5 (Figure 4, Panel E) highlighted biological processes related to “mitotic cell activity”, “flower” and “plant development”. These molecular mechanisms are potentially involved in the early bud break date observed in the population from low elevation and thus may explained their longest growing period. Genes related to “plant growth” were mainly found in cluster#1 (32 genes). Seven genes, for which we suggest a potential role in an early restart of meristem activity retain our attention: (i) a GR-RBP2 and 2 emb1138 genes encoding a glycine lycine-rich RNA-binding protein and DEAD box RNA helicases, respectively. members of these two gene families are known to be involved both in cold stress tolerance and in the acceleration of seed germination and seedling growth under low temperature (Kim *et al*., 2007; Gu *et al*., 2014), (ii) a FBL17 gene similar to an ubiquitin protein ligase FBL17 was also detected. Noir *et al*., (2015) showed that the loss of FBL17 function reduced drastically cell division in the apical meristem. Alike, a SHR gene similar to a GRAS family transcription factor, known to be a key regulator of cell division and meristem activity was also identified (Wang *et al*., 2011), (iii) a SE gene encoding a C2H2 zinc-finger protein Serrate (SE) protein known to accelerate leaf production and shorter time to flowering was also identified (Wilson *et al*., 2008), finally, (iv) we identified two genes (ARIA and F21P24.13) involved in ABA sensitivity, thermotolerance and seedling growth suggesting a potential role of these gene in the restart of meristem activity (Lee *et al*., 2010).

We also identified four and six genes related to the “flower development. Notably, cluster#4 included a LOX3 and a GA20×2 gene involved in hormone signaling. The LOX3 gene encode a lipoxygenase known to be probably involved in the biosynthesis of the jasmonic acid and influence flower development (Cao et al., 2016). The GA2OX 2 is similar to a Gibberellin 2-beta-dioxygenase involved in the gibberelin biosynthesis pathway. Yamauchi *et al*., (2007) reported that this gene partly suppresses the germination during dark exposition after inactivation of phytochrome. A F15K9.22 gene similar to a Fantastic Four (FAF) protein was also found in this cluster. This gene influence the activity of the shoot apical meristem and is upregulated once flowering is initiated (Wahl *et al*., 2010).

In cluster#5, we identified three genes (RAP2.7, GAI and LP1) for which we suggest a potential role both the early budburst and flowering date observed in populations from low elevation: (i) RAP2.7 similar to an Apetala2 protein. Okamuro *et al*., (1997) showed that this gene is active in the meristem both during reproductive and vegetative development, (ii) a GAI gene similar to a transcription factor involved in gibberellin signaling. In *Arabidopsis thaliana* upregulation of this gene accelerate flowering (Petty *et al*., 2003), and (iii) LP1 gene encoding a Non-specific lipid transfer protein. Using RNAi Arabidopsis line, Nieuwland *et al*., (2005) showed an important role of the LP1 gene in meristem functioning. Downregulation of the LP1 gene resulted in dwarfed plants and a disruption of flower development.

## Conclusion

In summary, our study provide a comprehensive overview of the sessile oak bud transcriptome along an elevation gradient. It reveals both a plastic response to temperature but also that this plastic response is genetically controlled, in that low and high altitude populations have evolved different molecular strategies to compromise a minimum of late frost damage and a maximum of vegetative period, thus increasing their respective fitness in such contrasted environmental conditions. Figure 5 graphically summarized our findings: the late budburst date observed for populations at high elevation was found to be related with an overexpression of genes involved in the inhibition of cell elongation and a delayed flowering time. Genes associated with cold tolerance were also found allowing buds to avoid late frost damage during early spring. Conversely, in populations at low elevation, we found a higher expression of genes involved in the acceleration of cell division and flowering, thus allowing buds to flush earlier when environmental once environmental conditions become favorable (Figure 5).

The molecular understanding of how locally adapted oak trees have fine-tuned their leaf unfolding process to a temperature gradient throughout the dormancy period, over micro-evolutionary time, provides valuable information to be further investigated in the light of the rise of temperature that accompany global warming.

## Supporting information

List of the primer pairs used for qPCR analysis.

qPCR validation of the selected candidate genes.

Overview of the sessile oak populations used in this study.

Evolution of phytohormone content in each population according to the Dormancy stage.

Overview of the cDNA libraries generated in this study.

Gene expression level comparison between the two valleys for a specific dormancy stage.

## Acknowledgements

This work was supported by the France Génomique (ANR-10-INBS-09-08, Oakadapt project). IL received funding from the European Research Council under the European Union’s Seventh Framework Program (FP/2007-2013) / ERC Treepeace Grant Agreement n. 339728. Illumina sequencing was performed at the Genoscope sequencing facility (www.genoscope.cns.fr). We thank the Genotoul Bioinformatics Platform Toulouse Occitanie (Bioinfo Genotoul, https://doi.org/10.15454/1.5572369328961167E12) for providing computing resources.

## Author contributions

GLP and CP designed the study. GLP, IL, and CP wrote the manuscript. GLP, CP and JML sampled the populations. GLP performed bioinformatics analysis with IL. GLP and CL were involved the RT-qPCR experiments. cDNA libraries construction and sequencing was done by KL, CDS and JMA. SD, AK and JML were involved in the monitoring of bud burst date and CP selected the populations for this study. All the authors read and approved the manuscript.

## Data accessibility

All the Illumina reads produced in this publication have been deposited in Short Read archive of NCBI. They can be recovered using the following study Accession number <u>PRJEB17876</u>.

## Supporting Information

**Supplementary Figure 1**: Evolution of phytohormone content in each population according to the Dormancy stage.

**Supplementary Figure 2**: qPCR validation of the selected candidate genes. **Supplementary Table 1**: Overview of the sessile oak populations used in this study.

**Supplementary Table 2**: List of the primer pairs used for qPCR analysis.

**Supplementary Table 3:** Overview of the cDNA libraries generated in this study.

**Supplementary Table 4**: Gene expression level comparison between the two valleys for a specific dormancy stage.

**Supplementary Files 1, 2, 3** are available online Inrae dataverse: G. Le Provost, C. Lalanne, I. Lesur’, JM. Louvet, S. Delzon, A. Kremer, K. Labadie, JM Aury, C Da Silva, T. Moritz, C Plomion,’ Locally adapted oak populations along an elevation gradient display different molecular strategies to regulate bud phenology, https://doi.org/10.15454/XMEKFX Portail Data INRAE, V3.0. These files includes normalized values for RNAseq data, Fold change Ratio, Gene set enrichment analysis and subnetwork enrichment analysis for genes displaying a significant dormancy, elevation and dormancy-by-elevation interaction effect respectively.

## References

Aitken SN, Yeaman S, Holliday JA, Wang T, Curtis-McLane S. 2008. Adaptation, migration or extirpation: climate change outcomes for tree populations. Evolutionary Applications 1: 95–111.

Alberto F, Bouffier L, Louvet J-M, Lamy J-B, Delzon S, Kremer A. 2011. Adaptive responses for seed and leaf phenology in natural populations of sessile oak along an altitudinal gradient. Journal of Evolutionary Biology 24: 1442–1454.

Alberto FJ, Derory J, Boury C, Frigerio J-M, Zimmermann NE, Kremer A. 2013. Imprints of natural selection along environmental gradients in phenology-related genes of Quercus petraea. Genetics 195: 495–512.

Alexa A. 2010. Gene set enrichment analysis with topGO. October: 27–27.

Anders S, Huber W. 2010. Differential expression analysis for sequence count data. Genome Biol 11: R106.

Ayala-Rodríguez JÁ, Barrera-Ortiz S, Ruiz-Herrera LF, López-Bucio J. 2017. Folic acid orchestrates root development linking cell elongation with auxin response and acts independently of the TARGET OF RAPAMYCIN signaling in Arabidopsis thaliana. Plant Science 264: 168–178.

Balandier P, Bonhomme M, Rageau R, Capitan F, Parisot E. 1993. Leaf bud endodormancy release in peach trees: evaluation of temperature models in temperate and tropical climates. Agricultural and Forest Meteorology 67: 95–113.

Bancos S, Nomura T, Sato T, Molnár G, Bishop GJ, Koncz C, Yokota T, Nagy F, Szekeres M. 2002. Regulation of transcript levels of the Arabidopsis cytochrome p450 genes involved in brassinosteroid biosynthesis. Plant Physiology 130: 504–513.

Björklund S, Antti H, Uddestrand I, Moritz T, Sundberg B. 2007. Cross-talk between gibberellin and auxin in development of Populus wood: gibberellin stimulates polar auxin transport and has a common transcriptome with auxin. The Plant Journal 52: 499–511.

Bouré N, Kumar SV, Arnaud N. 2019. The BAP Module: A Multisignal Integrator Orchestrating Growth. Trends in Plant Science 24: 602–610.

Bresson CC, Vitasse Y, Kremer A, Delzon S. 2011. To what extent is altitudinal variation of functional traits driven by genetic adaptation in European oak and beech? Tree Physiology 31: 1164–1174.

Burke JJ. 2001. Identification of genetic diversity and mutations in higher plant acquired thermotolerance. Physiologia Plantarum 112: 167–170.

Cabello JV, Arce AL, Chan RL. 2012. The homologous HD-Zip I transcription factors HaHB1 and AtHB13 confer cold tolerance via the induction of pathogenesis-related and glucanase proteins. The Plant Journal 69: 141–153.

Cao S, Chen H, Zhang C, Tang Y, Liu J, Qi H. 2016. Heterologous Expression and Biochemical Characterization of Two Lipoxygenases in Oriental Melon, Cucumis melo var. makuwa Makino. PLoS ONE 11.

Capblancq T, Morin X, Gueguen M, Renaud J, Lobreaux S, Bazin E. 2020. Climate-associated genetic variation in Fagus sylvatica and potential responses to climate change in the French Alps. Journal of Evolutionary Biology 33: 783–796.

Chao WS, Dogramaci M, Horvath DP, Anderson JV, Foley ME. 2017. Comparison of phytohormone levels and transcript profiles during seasonal dormancy transitions in underground adventitious buds of leafy spurge. Plant Molecular Biology 94: 281–302.

Cline MG, Deppong DO. 1999. The Role of Apical Dominance in Paradormancy of Temperate Woody Plants: A Reappraisal. Journal of Plant Physiology 155: 350–356.

Cooper RD, Shaffer HB. 2021. Allele-specific expression and gene regulation help explain transgressive thermal tolerance in non-native hybrids of the endangered California tiger salamander (Ambystoma californiense). Molecular Ecology 30: 987–1004.

Dantec CF, Vitasse Y, Bonhomme M, Louvet J-M, Kremer A, Delzon S. 2014. Chilling and heat requirements for leaf unfolding in European beech and sessile oak populations at the southern limit of their distribution range. International Journal of Biometeorology 58: 1853–1864.

DeYoung BJ, Bickle KL, Schrage KJ, Muskett P, Patel K, Clark SE. 2006. The CLAVATA1-related BAM1, BAM2 and BAM3 receptor kinase-like proteins are required for meristem function in Arabidopsis. The Plant Journal: For Cell and Molecular Biology 45: 1–16.

Dierck R, Keyser ED, Riek JD, Dhooghe E, Huylenbroeck JV, Prinsen E, Straeten DVD. 2016. Change in Auxin and Cytokinin Levels Coincides with Altered Expression of Branching Genes during Axillary Bud Outgrowth in Chrysanthemum. PLOS ONE 11: e0161732.

Doi H, Katano I. 2008. Phenological timings of leaf budburst with climate change in Japan. Agricultural and Forest Meteorology 148: 512–516.

Ducousso A, Guyon JP, Krémer A. 1996. Latitudinal and altitudinal variation of bud burst in western populations of sessile oak (Quercus petraea (Matt) Liebl). Annales des Sciences Forestières 53: 775– 782.

Fang S-C, Fernandez DE. 2002. Effect of Regulated Overexpression of the MADS Domain Factor AGL15 on Flower Senescence and Fruit Maturation. Plant Physiology 130: 78–89.

Firmat C, Delzon S, Louvet J-M, Parmentier J, Kremer A. 2017. Evolutionary dynamics of the leaf phenological cycle in an oak metapopulation along an elevation gradient. Journal of Evolutionary Biology 30: 2116–2131.

Fleet CM, Sun T. 2005. A DELLAcate balance: the role of gibberellin in plant morphogenesis. Current Opinion in Plant Biology 8: 77–85.

Footitt S, Müller K, Kermode AR, Finch-Savage WE. 2015. Seed dormancy cycling in Arabidopsis: chromatin remodelling and regulation of DOG1 in response to seasonal environmental signals. The Plant Journal 81: 413–425.

Fujino K, Matsuda Y. 2009. Genome-wide analysis of genes targeted by qLTG3-1 controlling low-temperature germinability in rice. Plant Molecular Biology 72: 137.

Gao R, Gruber MY, Amyot L, Hannoufa A. 2018. SPL13 regulates shoot branching and flowering time in Medicago sativa. Plant Molecular Biology 96: 119–133.

Gargul JM, Mibus H, Serek M. 2017. Characterization of Transgenic Kalanchoë and Petunia with Organ-Specific Expression of GUS or GA2ox Genes Led by the Deletion BOX-I Version (dBI) of the PAL1 Promoter. Journal of Plant Growth Regulation 36: 424–435.

Gu L, Xu T, Lee K, Lee KH, Kang H. 2014. A chloroplast-localized DEAD-box RNA helicaseAtRH3 is essential for intron splicing and plays an important role in the growth and stress response in Arabidopsis thaliana. Plant Physiology and Biochemistry 82: 309–318.

Hall D, Luquez V, Garcia VM, St Onge KR, Jansson S, Ingvarsson PK. 2007. Adaptive population differentiation in phenology across a latitudinal gradient in European aspen (Populus tremula, L.): a comparison of neutral markers, candidate genes and phenotypic traits. Evolution; International Journal of Organic Evolution 61: 2849–2860.

Hamant O, Pautot V. 2010. Plant development: A TALE story. Comptes Rendus Biologies 333: 371– 381.

He S-B, Wang W-X, Zhang J-Y, Xu F, Lian H-L, Li L, Yang H-Q. 2015. The CNT1 Domain of Arabidopsis CRY1 Alone Is Sufficient to Mediate Blue Light Inhibition of Hypocotyl Elongation. Molecular Plant 8: 822–825.

Heide OM. 2008. Interaction of photoperiod and temperature in the control of growth and dormancy of Prunus species. Scientia Horticulturae 115: 309–314.

Horvath D. 2010. Bud Dormancy and Growth. In: Pua EC, Davey MR, eds. Plant Developmental Biology - Biotechnological Perspectives. Berlin, Heidelberg: Springer Berlin Heidelberg, 53–70.

Horvath DP, Anderson JV, Chao WS, Foley ME. 2003. Knowing when to grow: signals regulating bud dormancy. Trends in Plant Science 8: 534–540.

Ingvarsson PK, García MV, Hall D, Luquez V, Jansson S. 2006. Clinal variation in phyB2, a candidate gene for day-length-induced growth cessation and bud set, across a latitudinal gradient in European aspen (Populus tremula). Genetics 172: 1845–1853.

Jue D, Sang X, Liu L, Shu B, Wang Y, Liu C, Xie J, Shi S. 2018. Identification of WRKY Gene Family from Dimocarpus longan and Its Expression Analysis during Flower Induction and Abiotic Stress Responses. International Journal of Molecular Sciences 19: 2169.

Karlgren A, Gyllenstrand N, Clapham D, Lagercrantz U. 2013. FLOWERING LOCUS T/TERMINAL FLOWER1-Like Genes Affect Growth Rhythm and Bud Set in Norway Spruce1[W][OPEN]. Plant Physiology 163: 792–803.

Kawecki TJ, Ebert D. 2004. Conceptual issues in local adaptation. Ecology Letters 7: 1225–1241.

Khalil-Ur-Rehman M, Sun L, Li C-X, Faheem M, Wang W, Tao J-M. 2017. Comparative RNA-seq based transcriptomic analysis of bud dormancy in grape. BMC Plant Biology 17: 18.

Kim HB, Lee H, Oh CJ, Lee H-Y, Eum HL, Kim H-S, Hong Y-P, Lee Y, Choe S, An CS, et al. 2010. Postembryonic Seedling Lethality in the Sterol-Deficient Arabidopsis cyp51A2 Mutant Is Partially Mediated by the Composite Action of Ethylene and Reactive Oxygen Species. Plant Physiology 152: 192–205.

Kim JY, Park SJ, Jang B, Jung C-H, Ahn SJ, Goh C-H, Cho K, Han O, Kang H. 2007. Functional characterization of a glycine-rich RNA-binding protein 2 in Arabidopsis thaliana under abiotic stress conditions. The Plant Journal 50: 439–451.

Kim SY, Shang Y, Joo S-H, Kim S-K, Nam KH. 2017. Overexpression of BAK1 causes salicylic acid accumulation and deregulation of cell death control genes. Biochemical and Biophysical Research Communications 484: 781–786.

Koehler G, Wilson RC, Goodpaster JV, Sønsteby A, Lai X, Witzmann FA, You J-S, Rohloff J, Randall SK, Alsheikh M. 2012. Proteomic Study of Low-Temperature Responses in Strawberry Cultivars (Fragaria × ananassa) That Differ in Cold Tolerance. Plant Physiology 159: 1787–1805.

Kremer A, Dupouey JL, Deans JD, Cottrell J, Csaikl U, Finkeldey R, Espinel S, Jensen J, Kleinschmit J, Van Dam B, et al. 2002. Leaf morphological differentiation between Quercus robur and Quercus petraea is stable across western European mixed oak stands. Annals of Forest Science 59: 777–787.

Kremer A, Potts BM, Delzon S. 2014. Genetic divergence in forest trees: understanding the consequences of climate change. Functional Ecology 28: 22–36.

Kremer A, Ronce O, Robledo-Arnuncio JJ, Guillaume F, Bohrer G, Nathan R, Bridle JR, Gomulkiewicz R, Klein EK, Ritland K, et al. 2012. Long-distance gene flow and adaptation of forest trees to rapid climate change. Ecology Letters 15: 378–392.

Lasky JR, Des Marais DL, Lowry DB, Povolotskaya I, McKay JK, Richards JH, Keitt TH, Juenger TE. 2014. Natural Variation in Abiotic Stress Responsive Gene Expression and Local Adaptation to Climate in Arabidopsis thaliana. Molecular Biology and Evolution 31: 2283–2296.

Le Provost G, Herrera R, Paiva JAP, Chaumeil P, Salin F, Plomion C. 2007. A micromethod for high throughput RNA extraction in forest trees. Biological Research 40: 291–297.

Le Provost G, Lesur I, Lalanne C, Da Silva C, Labadie K, Aury JM, Leple JC, Plomion C. 2016. Implication of the suberin pathway in adaptation to waterlogging and hypertrophied lenticels formation in pedunculate oak (Quercus robur L.) (C-J Tsai, Ed.). Tree Physiology: tpw056.

Le Provost G, Sulmon C, Frigerio JM, Bodénès C, Kremer A, Plomion C. 2012. Role of waterlogging-responsive genes in shaping interspecific differentiation between two sympatric oak species. Tree Physiology 32: 119–134.

Lee S, Cho D-I, Kang J, Kim M-D, Kim SY. 2010. AtNEK6 interacts with ARIA and is involved in ABA response during seed germination. Molecules and Cells 29: 559–566.

Lenoir J, Gégout JC, Marquet PA, Ruffray P de, Brisse H. 2008. A Significant Upward Shift in Plant Species Optimum Elevation During the 20th Century. Science 320: 1768–1771.

Lenoir J, Svenning J-C. 2015. Climate-related range shifts – a global multidimensional synthesis and new research directions. Ecography 38: 15–28.

Leroy T, Louvet J-M, Lalanne C, Provost GL, Labadie K, Aury J-M, Delzon S, Plomion C, Kremer A. 2020. Adaptive introgression as a driver of local adaptation to climate in European white oaks. New Phytologist 226: 1171–1182.

Lesur I, Bechade A, Klopp C, Noirot C, Leplé JC, Kremer A, Plomion C, Le Provost G. 2015a. A unigene set for European beech (Fagus sylvatica L.) and its use to decipher the molecular mechanisms involved in dormancy regulation. Molecular Ecology Ressources.

Lesur I, Provost GL, Bento P, Silva CD, Leplé J-C, Murat F, Ueno S, Bartholomé J, Lalanne C, Ehrenmann F, et al. 2015b. The oak gene expression atlas: insights into Fagaceae genome evolution and the discovery of genes regulated during bud dormancy release. BMC Genomics 16: 112.

Li W-F, Ding Q, Cui K-M, He X-Q. 2013. Cambium reactivation independent of bud unfolding involves de novo IAA biosynthesis in cambium regions in Populus tomentosa Carr. Acta Physiologiae Plantarum 35: 1827–1836.

Li F, Wu X, Lam P, Bird D, Zheng H, Samuels L, Jetter R, Kunst L. 2008. Identification of the Wax Ester Synthase/Acyl-Coenzyme A:Diacylglycerol Acyltransferase WSD1 Required for Stem Wax Ester Biosynthesis in Arabidopsis. Plant Physiology 148: 97–107.

Li J, Xu Y, Niu Q, He L, Teng Y, Bai S. 2018. Abscisic Acid (ABA) Promotes the Induction and Maintenance of Pear (Pyrus pyrifolia White Pear Group) Flower Bud Endodormancy. International Journal of Molecular Sciences 19.

Liu Y, Li X, Ma D, Chen Z, Wang J-W, Liu H. 2018. CIB1 and CO interact to mediate CRY2-dependent regulation of flowering. EMBO reports 19: e45762.

Lobréaux S, Miquel C. 2020. Identification of Arabis alpina genomic regions associated with climatic variables along an elevation gradient through whole genome scan. Genomics 112: 729–735.

Luedeling E, Girvetz EH, Semenov MA, Brown PH. 2011. Climate Change Affects Winter Chill for Temperate Fruit and Nut Trees. PLOS ONE 6: e20155.

Maurya JP, Bhalerao RP. 2017. Photoperiod- and temperature-mediated control of growth cessation and dormancy in trees: a molecular perspective. Annals of Botany 120: 351–360.

Mead A, Ramirez JP, Bartlett MK, Wright JW, Sack L, Sork VL. 2019. Seedling response to water stress in valley oak (Quercus lobata) is shaped by different gene networks across populations. Molecular Ecology 28: 5248–5264.

Menzel A. 2002. Phenology: Its Importance to the Global Change Community. Climatic Change 54: 379–385.

Min Z, Zhao X, Li R, Yang B, Liu M, Fang Y. 2017. Comparative transcriptome analysis provides insight into differentially expressed genes related to bud dormancy in grapevine (Vitis vinifera). Scientia Horticulturae 225: 213–220.

Moschen S, Bengoa Luoni S, Paniego NB, Hopp HE, Dosio GAA, Fernandez P, Heinz RA. 2014. Identification of Candidate Genes Associated with Leaf Senescence in Cultivated Sunflower (Helianthus annuus L.). PLoS ONE 9.

Naor A, Flaishman M, Stern R, Moshe A, Erez A. 2003. Temperature Effects on Dormancy Completion of Vegetative Buds in Apple. Journal of the American Society for Horticultural Science 128: 636–641.

Nieuwland J, Feron R, Huisman BAH, Fasolino A, Hilbers CW, Derksen J, Mariani C. 2005. Lipid Transfer Proteins Enhance Cell Wall Extension in Tobacco. The Plant Cell 17: 2009–2019.

Noir S, Marrocco K, Masoud K, Thomann A, Gusti A, Bitrian M, Schnittger A, Genschik P. 2015. The Control of Arabidopsis thaliana Growth by Cell Proliferation and Endoreplication Requires the F-Box Protein FBL17. The Plant Cell 27: 1461–1476.

Okamuro JK, Caster B, Villarroel R, Montagu MV, Jofuku KD. 1997. The AP2 domain of APETALA2 defines a large new family of DNA binding proteins in Arabidopsis. Proceedings of the National Academy of Sciences 94: 7076–7081.

Olsen JE, Lee Y, Junttila O. 2014. Effect of alternating day and night temperature on short day-induced bud set and subsequent bud burst in long days in Norway spruce. Frontiers in Plant Science 5.

Paul A, Jha A, Bhardwaj S, Singh S, Shankar R, Kumar S. 2014. RNA-seq-mediated transcriptome analysis of actively growing and winter dormant shoots identifies non-deciduous habit of evergreen tree tea during winters. Scientific Reports 4: 5932.

Petty LM, Harberd NP, Carré IA, Thomas B, Jackson SD. 2003. Expression of the Arabidopsis gai gene under its own promoter causes a reduction in plant height in chrysanthemum by attenuation of the gibberellin response. Plant Science 164: 175–182.

Plomion C, Aury J-M, Amselem J, Leroy T, Murat F, Duplessis S, Faye S, Francillonne N, Labadie K, Provost GL, et al. 2018. Oak genome reveals facets of long lifespan. Nature Plants: 1.

Racolta A, Nodine MD, Davies K, Lee C, Rowe S, Velazco Y, Wellington R, Tax FE. 2018. A Common Pathway of Root Growth Control and Response to CLE Peptides Through Two Receptor Kinases in Arabidopsis. Genetics 208: 687–704.

Rellstab C, Zoller S, Walthert L, Lesur I, Pluess AR, Graf R, Bodénès C, Sperisen C, Kremer A, Gugerli F. 2016. Signatures of local adaptation in candidate genes of oaks (Quercus spp.) with respect to present and future climatic conditions. Molecular Ecology 25: 5907–5924.

Ren H, Gray WM. 2015. SAUR Proteins as Effectors of Hormonal and Environmental Signals in Plant Growth. Molecular Plant 8: 1153–1164.

Rohde A, Bhalerao RP. 2007. Plant dormancy in the perennial context. Trends in Plant Science 12: 217–223.

Rozen S, Skaletsky H. 2000. Primer3 on the WWW for general users and for biologist programmers. Methods in molecular biology (Clifton, N.J.) 132: 365–386.

Ruttink T, Arend M, Morreel K, Storme V, Rombauts S, Fromm J, Bhalerao RP, Boerjan W, Rohde A. 2007. A molecular timetable for apical bud formation and dormancy induction in poplar. The Plant Cell 19: 2370–2390.

Santamaría ME, Rodríguez R, Cañal MJ, Toorop PE. 2011. Transcriptome analysis of chestnut (Castanea sativa) tree buds suggests a putative role for epigenetic control of bud dormancy. Annals of Botany 108: 485–498.

Savolainen O, Pyhäjärvi T, Knürr T. 2007. Gene Flow and Local Adaptation in Trees. Annual Review of Ecology, Evolution, and Systematics 38: 595–619.

Schaber J, Badeck F-W. 2003. Physiology-based phenology models for forest tree species in Germany. International Journal of Biometeorology 47: 193–201.

Sharan R, Maron-Katz A, Shamir R. 2003. CLICK and EXPANDER: a system for clustering and visualizing gene expression data. Bioinformatics (Oxford, England) 19: 1787–1799.

Sharma M, Banday ZZ, Shukla BN, Laxmi A. 2019. Glucose-Regulated HLP1 Acts as a Key Molecule in Governing Thermomemory. Plant Physiology 180: 1081–1100.

Shim D, Ko J-H, Kim W-C, Wang Q, Keathley DE, Han K-H. 2014. A molecular framework for seasonal growth-dormancy regulation in perennial plants. Horticulture Research 1: 14059.

Sork VL. 2018. Genomic Studies of Local Adaptation in Natural Plant Populations. Journal of Heredity 109: 3–15.

Sork VL, Squire K, Gugger PF, Steele SE, Levy ED, Eckert AJ. 2016. Landscape genomic analysis of candidate genes for climate adaptation in a California endemic oak, Quercus lobata. American Journal of Botany 103: 33–46.

Stief A, Altmann S, Hoffmann K, Pant BD, Scheible W-R, Bäurle I. 2014. Arabidopsis miR156 Regulates Tolerance to Recurring Environmental Stress through SPL Transcription Factors. The Plant Cell 26: 1792–1807.

Takemura Y, Kuroki K, Shida Y, Araki S, Takeuchi Y, Tanaka K, Ishige T, Yajima S, Tamura F. 2015. Comparative Transcriptome Analysis of the Less-Dormant Taiwanese Pear and the Dormant Japanese Pear during Winter Season. PLOS ONE 10: e0139595.

Tan H, Man C, Xie Y, Yan J, Chu J, Huang J. 2019. A Crucial Role of GA-Regulated Flavonol Biosynthesis in Root Growth of Arabidopsis. Molecular Plant 12: 521–537.

Tanino KK, Kalcsits L, Silim S, Kendall E, Gray GR. 2010. Temperature-driven plasticity in growth cessation and dormancy development in deciduous woody plants: a working hypothesis suggesting how molecular and cellular function is affected by temperature during dormancy induction. Plant Molecular Biology 73: 49–65.

Tavazoie S, Hughes JD, Campbell MJ, Cho RJ, Church GM. 1999. Systematic determination of genetic network architecture. Nature Genetics 22: 281–285.

Tylianakis JM, Didham RK, Bascompte J, Wardle DA. 2018. Global change and species interactions in terrestrial ecosystems. Ecology Letters: 1351–1363.

Ueno S, Klopp C, Leplé JC, Derory J, Noirot C, Léger V, Prince E, Kremer A, Plomion C, Provost GL. 2013. Transcriptional profiling of bud dormancy induction and release in oak by next-generation sequencing. BMC Genomics 14: 236.

Vitasse Y, Delzon S, Bresson CC, Michalet R, Kremer A. 2009a. Altitudinal differentiation in growth and phenology among populations of temperate-zone tree species growing in a common garden. Canadian journal of forest research.

Vitasse Y, Delzon S, Dufrêne E, Pontailler J-Y, Louvet J-M, Kremer A, Michalet R. 2009b. Leaf phenology sensitivity to temperature in European trees: Do within-species populations exhibit similar responses? Agricultural and Forest Meteorology 149: 735–744.

Vitasse Y, Porté AJ, Kremer A, Michalet R, Delzon S. 2009c. Responses of canopy duration to temperature changes in four temperate tree species: relative contributions of spring and autumn leaf phenology. Oecologia 161: 187–198.

Wang J, Andersson-Gunnerås S, Gaboreanu I, Hertzberg M, Tucker MR, Zheng B, Leśniewska J, Mellerowicz EJ, Laux T, Sandberg G, et al. 2011. Reduced Expression of the SHORT-ROOT Gene Increases the Rates of Growth and Development in Hybrid Poplar and Arabidopsis. PLoS ONE 6.

Wang L-J, Li S-H. 2006. Salicylic acid-induced heat or cold tolerance in relation to Ca2+ homeostasis and antioxidant systems in young grape plants. Plant Science 170: 685–694.

Wang Y, Li L, Ye T, Lu Y, Chen X, Wu Y. 2013a. The inhibitory effect of ABA on floral transition is mediated by ABI5 in Arabidopsis. Journal of Experimental Botany 64: 675–684.

Wang C, Yan X, Chen Q, Jiang N, Fu W, Ma B, Liu J, Li C, Bednarek SY, Pan J. 2013b. Clathrin Light Chains Regulate Clathrin-Mediated Trafficking, Auxin Signaling, and Development in Arabidopsis. The Plant Cell 25: 499–516.

White TL, Adams WT, Neale DB. 2007. Genetic markers - morphological, biochemical and molecular markers. Forest genetics: 53–74.

Wilson MD, Wang D, Wagner R, Breyssens H, Gertsenstein M, Lobe C, Lu X, Nagy A, Burke RD, Koop BF, et al. 2008. ARS2 Is a Conserved Eukaryotic Gene Essential for Early Mammalian Development. Molecular and Cellular Biology 28: 1503–1514.

Yamauchi Y, Takeda-Kamiya N, Hanada A, Ogawa M, Kuwahara A, Seo M, Kamiya Y, Yamaguchi S. 2007. Contribution of Gibberellin Deactivation by AtGA2ox2 to the Suppression of Germination of Dark-Imbibed Arabidopsis thaliana Seeds. Plant and Cell Physiology 48: 555–561.

Yang Q, Gao Y, Wu X, Moriguchi T, Bai S, Teng Y. 2021. Bud endodormancy in deciduous fruit trees: advances and prospects. Horticulture Research 8: 1–11.

Yang Q-S, Wu J-H, Li C-Y, Wei Y-R, Sheng O, Hu C-H, Kuang R-B, Huang Y-H, Peng X-X, McCardle JA, et al. 2012a. Quantitative Proteomic Analysis Reveals that Antioxidation Mechanisms Contribute to Cold Tolerance in Plantain (Musa paradisiaca L.; ABB Group) Seedlings. Molecular & Cellular Proteomics 11: 1853–1869.

Yang X, Yu Y, Jiang L, Lin X, Zhang C, Ou X, Osabe K, Liu B. 2012b. Changes in DNA methylation and transgenerational mobilization of a transposable element (mPing) by the Topoisomerase II inhibitor, Etoposide, in rice. BMC Plant Biology 12: 48.

Yordanov YS, Ma C, Strauss SH, Busov VB. 2014. EARLY BUD-BREAK 1 (EBB1) is a regulator of release from seasonal dormancy in poplar trees. Proceedings of the National Academy of Sciences: 201405621.

Zanetto A, Kremer A. 1995. Geographical structure of gene diversity in Quercus petraea (Matt.) Liebl. I. Monolocus patterns of variation. Heredity 75: 506–517.

Zhao Y, Cheng S, Song Y, Huang Y, Zhou S, Liu X, Zhou D-X. 2015. The Interaction between Rice ERF3 and WOX11 Promotes Crown Root Development by Regulating Gene Expression Involved in Cytokinin Signaling. The Plant Cell 27: 2469–2483.

Zhong W, Gao Z, Zhuang W, Shi T, Zhang Z, Ni Z. 2013. Genome-wide expression profiles of seasonal bud dormancy at four critical stages in Japanese apricot. Plant Molecular Biology 83: 247–264.

Zhu Y, Shi Z, Li S, Liu H, Liu F, Niu Q, Li C, Wang J, Rong T, Yi H, et al. 2018. Fine mapping of the novel male-sterile mutant gene ms39 in maize originated from outer space flight. Molecular Breeding 38: 125.

