## Supplementary material for "Locally adapted oak populations along an elevation gradient display different molecular strategies to regulate bud phenology": List of the primer pairs used for qPCR analysis.

**Supplementary Table 2**: List of the primer pairs used for qPCR analysis. Abbreviations: Tm: annealing temperature, E: Elevation effect, D*E: Dormancy-by-elevation effect. For each effect, the cluster ID is indicated in parenthesis in the second column. Qrob_IDs were retrieved from the oak genome available in Plomion et al. (2018).

.

| Gene ID | Significant effect in  RNAseq | Function | Forward primer(5’3’)  Reverse primer (3’-5’) | Amplicon  size (bp) | Multiband in  agarose gel | Used in  qPCR | PCR  Efficiency | Highest expression in qPCR |
| --- | --- | --- | --- | --- | --- | --- | --- | --- |
| Qrob_P0007950.2 | E (CL1) | **Disease resistance-responsive protein** | GCAATCCCTCTTCAGTCCCA  CACACTGAGTTCCTGCCTAG | 276 | No | Yes | 103 | **High elevation** |
| Qrob_P0292970.2 | E (CL1) | **Unknown protein** | TGTTGTGATGAAGGCAGACG  ACAGCAACTCCCTCATCCAA | 192 | No | Yes | 108 | **High elevation** |
| Qrob_P0134460.2 | E (CL4) | **Metal transport/detoxification superfamily protein** | GTCCAAGGCCATGCAGATTG  GTACTGACTGGCGAGACACT | 177 | No | Yes | 95 | **High elevation** |
| Qrob_P0247760.2 | E(CL4) | **TIR-NBS-LRR** | GACGGCTTTCATGACCAACA  AATTTCTTCTCCCCTCGGCA | 181 | Yes | No | NA | **NA** |
| Qrob_P0059940.2 | E(CL2) | **G-type lectin S-receptor-like serine/threonine-protein kinase** | GACACCATCTCTGCACACCA GTCTCTCTGTTTGCCACCCA | 179 | No | yes | 97 | **Low elevation** |
| Qrob_P0201490.2 | E(CL5) | **Urine biosynthesis 4 protein** | TTTTCCTCGCCCCAATAGGT  ATAGGGTGCGTCAGTTGGAA | 244 | No | yes | 107 | **Low/Mean**  **elevation** |
| Qrob_P0440000.2 | D*E (CL2) | **Sucrose synthase 3** | TTTTCCTCGCCCCAATAGGT  ATAGGGTGCGTCAGTTGGAA | 214 | No | yes | 103 | **D*E** |
| Qrob_P0477210.2 | D*E (CL2) | **Pyrophosphorylase 2** | TGATCCTGAGTTCCGCCATT  GGCTTCAATGGCAGACTCAG | 151 | No | Yes | 98 | **D*E** |
| Qrob_P0252420.2 | D*E (CL3) | **Amino Acid permease 7** | ACCTGATGAACCTGAGGAGC  TTCTAGCAGGGCCACATTCA | 236 | Yes | No | NA | **NA** |
| Qrob_P0140130.2 | D*E (CL4) | **TCP domain protein 9** | GATGCAAGTGACGCCAGTAG  AGTTCTCGGGTCAGCTGAAA | 171 | No | Yes | 98 | **D*E** |
| Qrob_P0768730.2 | D*E (CL4) | **GDP-D-mannose 4,6-dehydratase 1** | AATTCAAGTGCTCCTCCGGT  GCTCTGCTTGTATCTCCCTT | 193 | No | Yes | 98 | **D*E** |
| Qrob_P0167140.2 | D*E (CL5) | **SAG101; Senescence Associated Gene 101** | GGCAAATTTCGTGGTGACCC TTGCTACTAGAGGGCCAGGT | 171 | No | Yes | 99 | **D*E** |
| **Control Genes** | | | | | | | | |
| Qrob_P0530610 | NS | **Unknown protein** | GAAGCACCACCCTCACAAGT  GTCTCCTCACAACTCACCGG | 197 | No | Yes | 110 | **NA** |
| Qrob_P0426000 | NS | **Unknown protein** | GCAGAGCTCCAGGACATGATaCAGCAGCAGAGATGAACCCA | 175 | No | Yes | 106 | **NA** |
