## Supplementary material for "Locally adapted oak populations along an elevation gradient display different molecular strategies to regulate bud phenology": qPCR validation of the selected candidate genes.

Supplementary Figure 2: qPCR validation. Panel (A) genes displaying a significant elevation effect. Panel (B) genes displaying a significant Dormancy-by-elevation effect. Abbreviations correspond to EndoD: Endodormant buds , EcoD: Ecodormant buds, Low: 100 mts (i.e. O-01+L-01), Mean: 800 mts (i.e. O-08+L-08) and High: 1,600 mts (i.e. O-16+L-16). Standard deviations were obtained from the four biological replicates

(A)

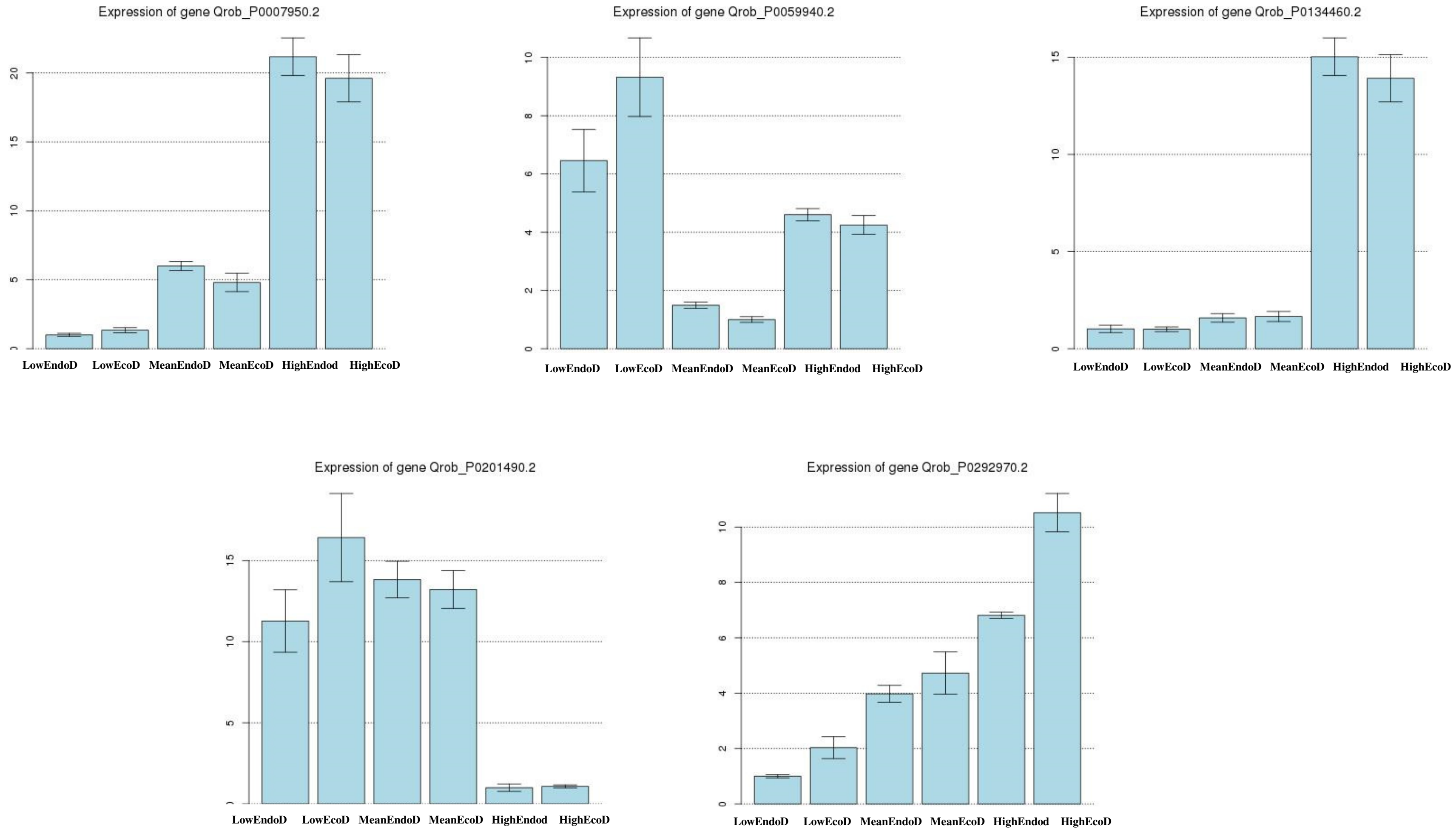

(B)

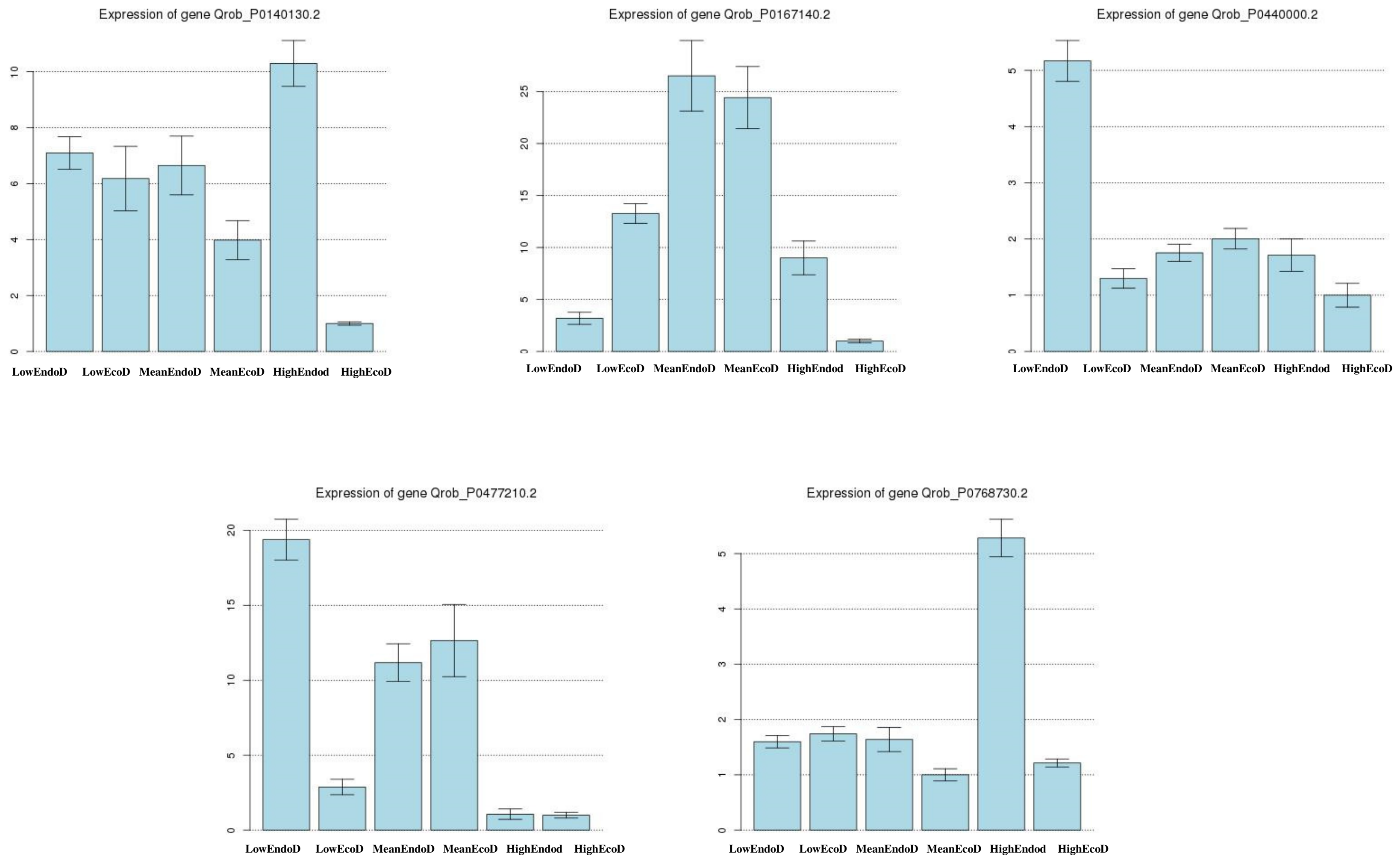
Expression of gene Qrob_P0140130.2
LowEndoD LowEcoD MeanEndoD MeanEcoD HighEndod HighEcoD
Expression of gene Qrob_P0167140.2
LowEndoD LowEcoD MeanEndoD MeanEcoD HighEndod HighEcoD
Expression of gene Qrob_P0440000.2
LowEndoD LowEcoD MeanEndoD MeanEcoD HighEndod HighEcoD
Expression of gene Qrob_P0477210.2
LowEndoD LowEcoD MeanEndoD MeanEcoD HighEndod HighEcoD
Expression of gene Qrob_P0768730.2
LowEndoD LowEcoD MeanEndoD MeanEcoD HighEndod HighEcoD
