## Supplementary material for "Locally adapted oak populations along an elevation gradient display different molecular strategies to regulate bud phenology": Overview of the sessile oak populations used in this study.

Supplementary Table 1: Overview of the sessile oak populations used in this study.

| **Populations** | **Valleys** | **Elevations (meters)** | **Latitude**  **Longitude** | **Endodormancy sampling dates** | **Ecodormancy sampling dates** |
| --- | --- | --- | --- | --- | --- |
| Laveyron  (O-01) | Ossau | 100 | 43°45’49” N  0° 13’11” W | 6 October 2013 | 12 March 2014 |
| Papillon  (O-08) | Ossau | 800 | 43° 25’ 23” N  0° 1’59” W | 7 October 2013 | 17 March 2014 |
| Péguere  (O16) | Ossau | 1,600 | 43° 52’ 00” N  0° 07’09” W | 8 October 2013 | 7 April 2014 |
| Josbaig  (L-01) | Luz | 100 | 42°13’35” N  0° 44’28” W | 10 October 2013 | 10 March 2014 |
| LeHourcq  (L-08) | Luz | 800 | 42°54’46” N  0° 26’04” W | 10 October 2013 | 16 March 2014 |
| Artouste  (L16) | Luz | 1,600 | 42°53’00” N  0° 24’08” W | 9 October 2013 | 7 April 2014 |
