## Supplementary material for "Locally adapted oak populations along an elevation gradient display different molecular strategies to regulate bud phenology": Evolution of phytohormone content in each population according to the Dormancy stage.

Supplementary Figure 1: Evolution phytohormone content in each population according to the Dormancy stage. Panel A: Table for ANOVA results for phytohormonre analysis. P value is indicated in each cell. *Pvalue<0.05, **Pvalue<0.01 and ***Pvalue<0.0001. NS stands for not significant. Panel B: Graphical representation of their accumulation over the Dormancy period. We used blue and orange color for endodormancy (i.e.Endo) and Ecodormancy (i.e. Eco) samples, respectivelly. Standard deviations were obtained from the 3 measurements performed in each population (Low=O-01+L-01, Mean=O-08+L-08 and High=O-18+L-18). Effects identified in the linear model where also indicated. Abbreviations correspond to: D: Dormancy effect, E: Elevation effect, D*E: interaction effect. * Pvalue<0.05, **Pvalue<0.001 and ***Pvalue<0.0001).

(A)

|  | IAA | ABA | Cytokinines |
| --- | --- | --- | --- |
| Dormancy | *******  **2.92 10^-6^** | *******  **0.000015** | ******  **0.00439** |
| Valley | NS  0.640 | NS  0.653 | NS  0.253 |
| Elevation | NS  0.735 | ******  **0.005** | NS  0.174 |

(B)

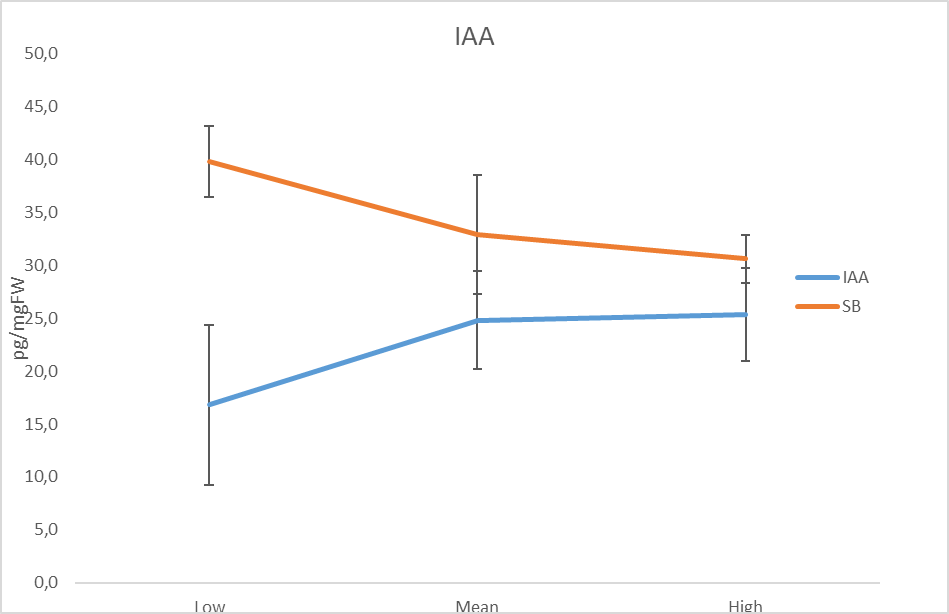

Endo

Eco

**D^*^ , E^***^ and D*E^***^**

IAA

pg/mgFW

|  | IAA | SB |
| --- | --- | --- |
| Low | 16.8267 | 39.7867 |
| Mean | 24.8167 | 32.925 |
| High | 25.3483 | 30.6167 |

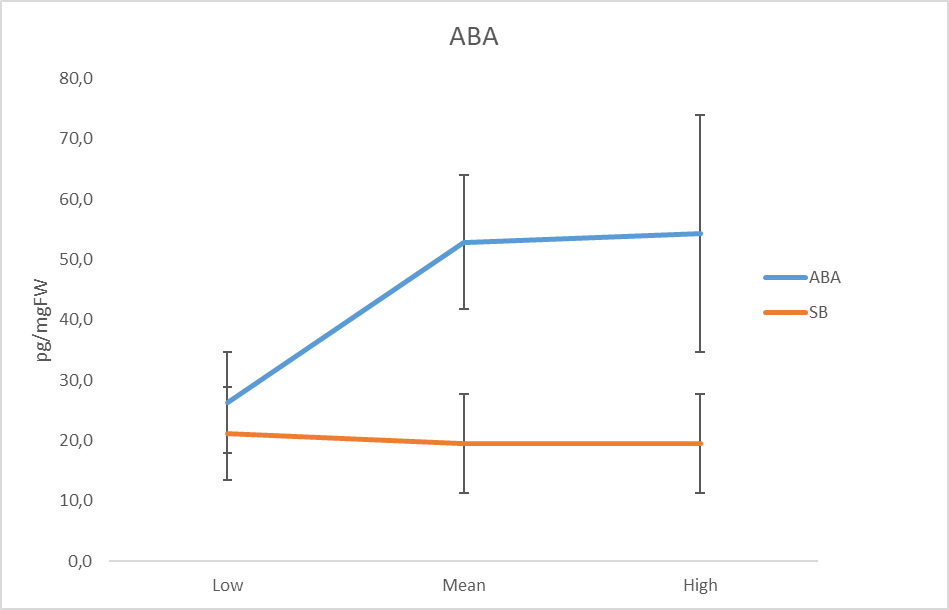

Endo

Eco

**D^***^** and **D*E^**^**

ABA

pg/mgFW

|  | ABA | SB |
| --- | --- | --- |
| Low | 26.3167 | 21.15 |
| Mean | 52.8333 | 19.5 |
| High | 54.2333 | 19.5 |

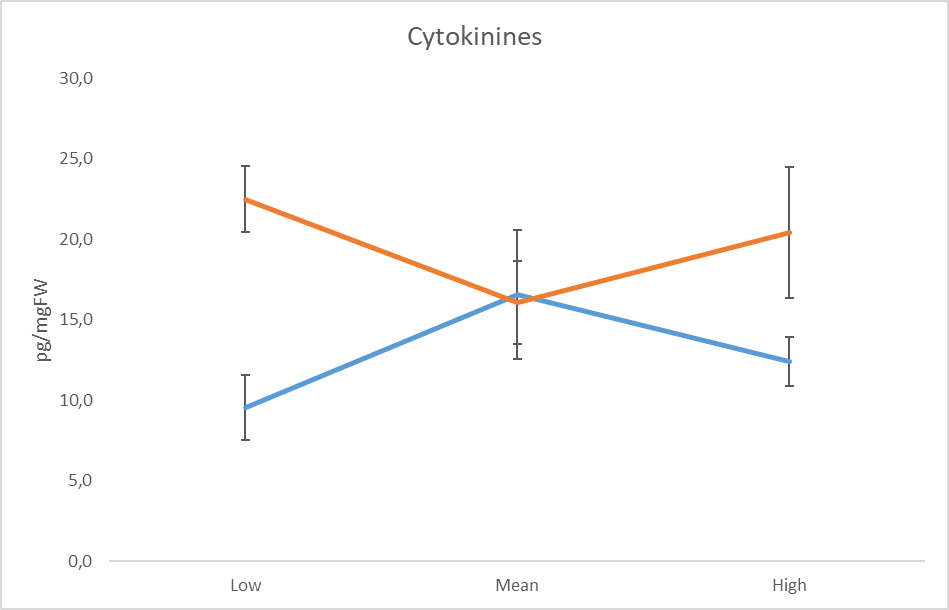

**D^***^**

Endo

Eco

Cytokinines

pg/mgFW

|  | Cytokinine | SB |
| --- | --- | --- |
| Low | 9.555 | 22.475 |
| Mean | 16.56 | 16.07 |
| High | 12.405 | 20.4 |

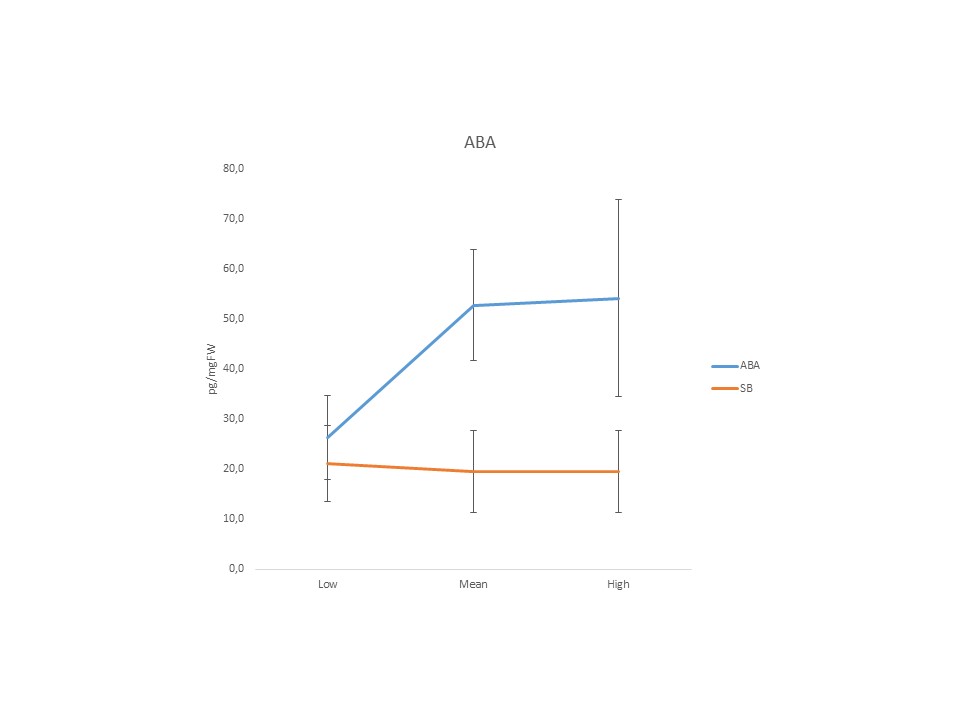
