## Supplementary material for "Locally adapted oak populations along an elevation gradient display different molecular strategies to regulate bud phenology": Overview of the cDNA libraries generated in this study.

Supplementary Table 3: Overview of the cDNA libraries generated in this study. Abbreviations correspond to: Endo for Endodormancy and Eco for Ecodormancy.

| **Library** | **Dormancy stage** | **Valley** | **Elevation in meters** | **Biological replicate** | **Number of read** | **Number of read mapped** | **% of mapped read** | **ENA study Accesion** |
| --- | --- | --- | --- | --- | --- | --- | --- | --- |
| **Josbaig 1 End (L-01)o** | **Endo** | **Luz** | **100** | **Bio. Rep. 1** | 33,041,456 | 26,108,308 | 78% | [PRJEB17876](https://www.ebi.ac.uk/ena/browser/view/PRJEB17876) |
| **Josbaig 2 Endo (L-01)** | **Endo** | **Luz** | **100** | **Bio. Rep. 2** | 39,872,577 | 30,912,308 | 76% |  |
| **Josbaig 1 Eco (L-01)** | **Eco** | **Luz** | **100** | **Bio. Rep. 1** | 50,981,819 | 41,289,518 | 81% |  |
| **Josbaig 2 Eco (L-01)** | **Eco** | **Luz** | **100** | **Bio. Rep. 2** | 23,561,827 | 19,087,016 | 82% |  |
| **Le Hourque 1 Endo**  **(L-08)** | **Endo** | **Luz** | **800** | **Bio. Rep. 1** | 30,616,326 | 23,634,328 | 76% |  |
| **Le Hourque 2 Endo**  **(L-08)** | **Endo** | **Luz** | **800** | **Bio. Rep. 2** | 29,858,750 | 23,458,930 | 79% |  |
| **Le Hourque 1 Eco**  **(L-08)** | **Eco** | **Luz** | **800** | **Bio. Rep. 1** | 27,938,013 | 21,945,696 | 77% |  |
| **Le Hourque 2 Eco**  **(L-08)** | **Eco** | **Luz** | **800** | **Bio. Rep. 2** | 26,431,779 | 21,060,600 | 80% |  |
| **Papillon 1 Endo**  **(O-08)** | **Endo** | **Ossau** | **800** | **Bio. Rep. 1** | 28,255,115 | 22,242,856 | 78% |  |
| **Papillon 2 Endo**  **(O-08)** | **Endo** | **Ossau** | **800** | **Bio. Rep. 2** | 28,507 89 | 22,407,716 | 78% |  |
| **Papillon 1 Eco**  **(O-08)** | **Eco** | **Ossau** | **800** | **Bio. Rep. 1** | 24,630,650 | 16,888,852 | 66% |  |
| **Papillon 2 Eco**  **(O-08)** | **Eco** | **Ossau** | **800** | **Bio. Rep. 2** | 30,949,276 | 22,751,666 | 73% |  |
| **Peguere 1 Endo**  **(O-16)** | **Endo** | **Ossau** | **1,600** | **Bio. Rep. 1** | 38,381,974 | 28,196,854 | 73% |  |
| **Peguere 2 Endo**  **(O-16)** | **Endo** | **Ossau** | **1,600** | **Bio. Rep. 2** | 33,474,713 | 25,096,294 | 75% |  |
| **Peguere 1 Eco**  **(O-16)** | **Eco** | **Ossau** | **1,600** | **Bio. Rep. 1** | 35,260,832 | 25,899,890 | 71% |  |
| **Peguere 2 Eco**  **(O-16)** | **Eco** | **Ossau** | **1,600** | **Bio. Rep. 2** | 37,076,987 | 27,232,020 | 72% |  |
| **Artouste 1 Endo**  **(L16)** | **Endo** | **Luz** | **1,600** | **Bio. Rep. 1** | 33,963,824 | 26,114,606 | 78% |  |
| **Artouste 2 Endo**  **(L16)** | **Endo** | **Luz** | **1,600** | **Bio. Rep. 2** | 32,241,220 | 25,563,258 | 78% |  |
| **Artouste 1 Eco**  **(L16)** | **Eco** | **Luz** | **1,600** | **Bio. Rep. 1** | 28,924,891 | 22,514,446 | 78% |  |
| **Artouste 2 Eco**  **(L16)** | **Eco** | **Luz** | **1,600** | **Bio. Rep. 2** | 28,146,228 | 20,581,192 | 71% |  |
| **Laveyron 1 Endo**  **(O-01)** | **Endo** | **Ossau** | **100** | **Bio. Rep. 1** | 32,668,746 | 25,573,700 | 78% |  |
| **Laveyron 2 Endo**  **(O-01)** | **Endo** | **Ossau** | **100** | **Bio. Rep. 2** | 29,920,489 | 23,661,520 | 79% |  |
| **Laveyron 1 Eco**  **(O-01)** | **Eco** | **Ossau** | **100** | **Bio. Rep. 1** | 33,368,451 | 27,291,694 | 81% |  |
| **Laveyron 2 Eco**  **(O-01)** | **Eco** | **Ossau** | **100** | **Bio. Rep. 2** | 33,982,054 | 27,638,362 | 81% |  |
