## Supplementary material for "Locally adapted oak populations along an elevation gradient display different molecular strategies to regulate bud phenology": Gene expression level comparison between the two valleys for a specific dormancy stage.

Supplementary Table 4: Gene expression level comparison between the two valleys for a specific dormancy stage. The comparison was performed for the population harvested at a same elevation in the two valleys considered. In each cell, we indicated the number of differentially expressed genes; the percentage is indicated in parenthesis.

|  | Endodormancy | Ecodormancy |
| --- | --- | --- |
| O-01 vs. L-01 | 76  (0.6%) | 81  (0.6%) |
| O-08 vs. L-08 | 149  (1.2%) | 78  (0.6%) |
| O-16 vs. L-16 | 55  (0.4%) | 93  (0.7%) |
